# Virus-derived variation in diverse human genomes

**DOI:** 10.1101/2020.11.20.390880

**Authors:** Shohei Kojima, Anselmo Jiro Kamada, Nicholas F. Parrish

## Abstract

Acquisition of genetic material from viruses by their hosts can generate inter-host structural genome variation. We developed computational tools enabling us to study virus-derived structural variants (SVs) in population-scale whole genome sequencing (WGS) datasets and applied them to 3,332 humans. Although SVs had already been cataloged in these subjects, we found previously-overlooked virus-derived SVs. We detected somatic SVs present in the sequenced lymphoblastoid cell lines (LCLs) derived from squirrel monkey retrovirus (SMRV), human immunodeficiency virus 1 (HIV-1), and human T lymphotropic virus (HTLV-1); these variants are attributable to infection of LCLs or their progenitor cells and may impact gene expression results and the biosafety of experiments using these cells. In addition, we detected new heritable SVs derived from human herpesvirus 6 (HHV-6) and human endogenous retrovirus-K (HERV-K). We report the first solo-DR HHV-6 that likely to reflects rearrangement of a known full-length endogenous HHV-6. We used linkage disequilibrium between single nucleotide variants (SNVs) and variants in reads that align to HERV-K, which often cannot be mapped uniquely using conventional short-read sequencing analysis methods, to locate previously-unknown polymorphic HERV-K loci. Some of these loci are tightly linked to trait-associated SNVs, some are in complex genome regions inaccessible to prior methods, and some contain novel HERV-K haplotypes likely derived from gene conversion from an unknown source or introgression. These tools and results broaden our perspective on the coevolution between viruses and humans, including ongoing virus-to-human gene transfer contributing to genetic variation between humans.

## Introduction

Union of genomes from discrete biological entities is a major engine of genetic diversity. Fusion of gametes, each bearing a set of recombinant chromosomes, is the immediate source of the genetic material that uniquely identifies each human. Taking a wider viewpoint, much of human genome can be recognized to have been acquired from a non-human source. For example, about 2% of the genome of many living humans can be attributed to introgression from Neanderthals (1). Movement of genetic information between biological entities apart from sexual reproduction, known as horizontal gene transfers (HGT), has also occurred in the human lineage. Some HGT happened so long ago that it is difficult to accurately classify the entity contributing the horizontally-transferred sequences according to extant taxonomies. This case for the bacteria, acquired millennia ago, now represented as our mitochondrial genomes. Other HGT occurred more recently. For example, about 8% of human genetic material is derived from human endogenous retroviruses (HERV) that integrated into our ancestors’ germline and then developed an intracellular replication cycle; some HERVs integrated recently enough that they can be classified based on homology to extant exogenous retroviruses.

The most recent among these retroviral integrations, of a lysine tRNA-primed HERV (i.e. HERV-K) subgroup called HML-2, occurred less than a million years ago (2). A single HERV-K element showing insertional polymorphism in different humans was known at the time of completion of the draft human genome (3), but during the past 20 years, over 40 insertionally-polymorphic elements have been described (2, 4–9). In addition to retroviruses, sequences from ancient relatives of Borna disease virus, an RNA-only virus, were horizontally acquired in the haplorrhine lineage (10). Human herpesviruses 6A and 6B, double stranded DNA viruses, have also been horizontally transferred to some human genomes during the holocene (11–13). These observations show that viruses acquired during the lifespan of an individual organism, including humans, have sometimes contributed to the genetic material passed on to their offspring, seemingly in violation of Weismann’s proposed barrier between soma and germline (14). When these viral sequences are acquired, the resulting mutation would be classified as a structural variant, defined as a DNA rearrangement greater than 50 nucleotides in length. Structural variants in human genomes are increasingly characterized at population scale (15–17). In these studies, SVs caused by polymorphic insertion of mobile genetic elements classified as transposons (including Alu, LINE-1, and SVA) have been considered explicitly. On the other hand, structural variants derived from viruses, another potentially important class of mobile genetic elements, have yet to be analyzed comprehensively.

Here we designed and applied new tools to comprehensively assess virus-derived structural variants in short-read genome sequencing data at population scale. We are not the first to consider viral sequences present in shotgun WGS datasets, however others have done so under the assumption that viral reads reflect a somatically-acquired “virome,” similar to the bacterial microbiome (18, 19). To distinguish exogenous virus contamination from germline integration, we applied several criteria, including read depth relative to autosomal genes and patterns of linkage disequilibrium with SNVs. Although we used human WGS datasets which have already been deeply analyzed to establish global SV references (15, 17), we discovered previously-undescribed heritable SVs derived from virus-origin genetic material. We detect squirrel monkey retrovirus (SMRV), human immunodeficiency virus 1 (HIV-1), and human T lymphotropic virus 1 (HTLV-1) in LCLs widely distributed as reference materials for characterizing human genetic and phenotypic variation, raising both biosafety and reproducibility concerns. We developed a new approach to detect and map polymorphisms in HERV-K that allows us to infer polymorphisms at over 60 loci previously unknown to be polymorphic, including new loci associated with human phenotypes. We show that viruses contribute unexpectedly to human genome structural variation and describe new tools for analyzing these variants at increasing scales.

## Results

### Detection of virus-mapped reads from diverse human populations

To discover human structural variation derived from viral sequences, we analyzed WGS reads that failed to map to the reference human genome (GRCh38DH). We used 3,332 high-coverage WGS datasets from the 1,000 Genomes Project (1kGP) and the Human Genome Diversity Project (HGDP) (20, 21), all derived from lymphoblastoid cell lines (LCLs). Unmapped reads were re-mapped to reference virus genomes from NCBI (see methods). We focused on viruses with abundantly-mapped reads, requiring that 5% of a viral genome be covered at more than 2x read depth. Applying this filter, we detected 7 high-coverage viruses (Figure 1, Supplementary Figure 1). Next we checked the patterns of viral genome coverage using the plots automatically generated as an output from our tool. We detected reads mapping to Chum salmon reovirus in 2 datasets from individuals of South Asian ancestry; only the first 200-bp of the viral genome was covered by reads, which were abundant in these two datasets but absent in others (Supplementary Figure 2). We detected reads mapping to simian T-lymphotropic virus 1 (STLV-1) in 4 samples. STLV-1-mapped reads were only detected in the datasets in which HTLV-1-mapped reads were also found (see below), and the same reads mapped to both HTLV-1 and STLV-1 (Supplementary Figure 2). In contrast, reads from SMRV, HIV-1, HTLV-1, human herpesvirus 6A (HHV-6A) and human herpesvirus 6B (HHV-6B) were abundantly detected in at least one subject, potentially consistent with presence in the germline, and reads covered the entire viral genome.

**Figure 1.**
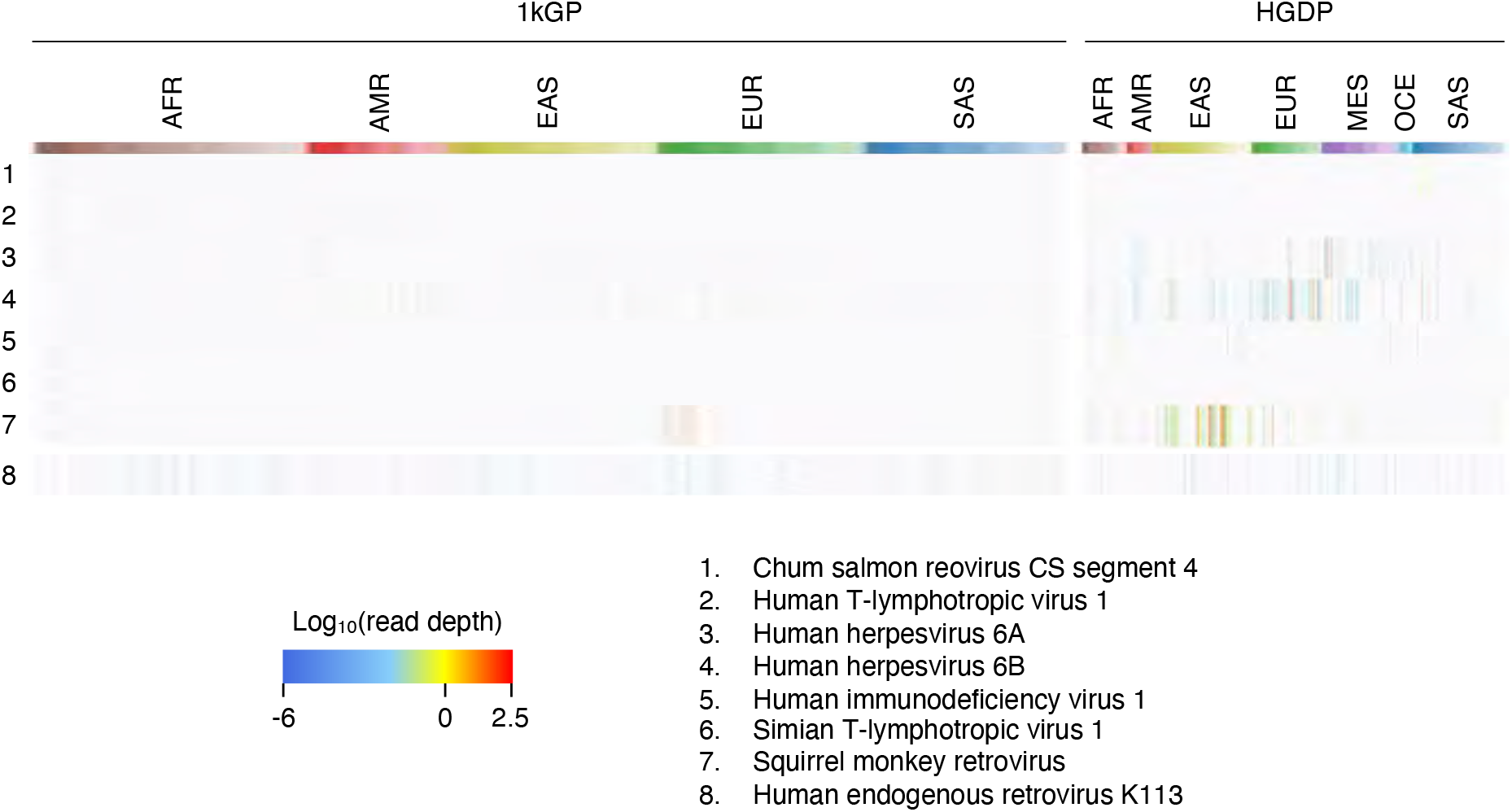
Virus search from 3,332 WGS. Heatmap shows read depth of seven viruses with abundant reads in at least one dataset and HERV-K113. The column colors show the human populations in the two databases. See Supplementary Figure 1 for the details of the names of the indicated populations. (1kGP: 1,000 Genomes Project; HGDP: Human Genome Diversity Project; AFR: African; AMR: American; EAS: East Asian; EUR: European; SAS: South Asian).

### Squirrel monkey retrovirus

SMRV-mapped reads were abundantly detected in 18 datasets, with a wide range of SMRV-mapped read depths in datasets from different subjects (Figure 2A, Supplementary Figure 3). From the 1kGP, all 12 datasets with SMRV were from subjects from Utah, including 2 subjects with read depths greater than 1x autosomal depth. All reads lack a guanosine in the tRNA primer binding site (PBS) relative to SMRV-H, which was isolated from macaque cell-derived preparations of Epstein Barr virus (EBV) (22); the tRNA PBS of the SMRV detected here is identical to SMRV sequences recently obtained from Vero cells (23), a cell line used for biologicals production (Figure 2A) (24). To determine if SMRV was integrated into the genomes of the sequenced LCLs, we searched for paired reads with one read mapped to the virus genome and one pair to the human genome (i.e. hybrid reads). Virus-human hybrid reads were observed, often mapped to the SMRV long terminal repeat (LTR), consistent with the virus being integrated into human chromosomes (Figure 2B); the human-mapped reads of these hybrid pairs mapped to multiple chromosomal loci (Figure 2C, Supplementary Figure 4). We found no enriched integration sites consistent with a clonal integration in the germline, nor shared integration sites across different subjects. These observations suggest these SMRV integrations are somatic rather than present in the germline. To assess whether the SMRV integrations occurred before or after the peripheral blood mononuclear cells (PBMCs) used to produce LCLs were removed from these subjects, we analyzed sequencing results obtained directly from these subject’s nucleated blood cells (25, 26). No reads mapped to SMRV. The most plausible source of SMRV in the sequenced cell lines is thus laboratory contamination (22, 27). This observation notwithstanding, SMRV requires biosafety level 2 (BSL2) containment, and the adventitious presence of SMRV in these samples could influence other results using these reference materials. We obtained LCLs from 2 subjects from whom high SMRV reads were found and confirmed the presence of SMRV DNA by PCR (Fig 2D). We analyzed existing RNAseq data (28) and confirmed that SMRV is transcribed and is associated with differential gene expression relative to LCLs in which SMRV RNA is not detected (Supplementary Figure 5).

**Figure 2.**
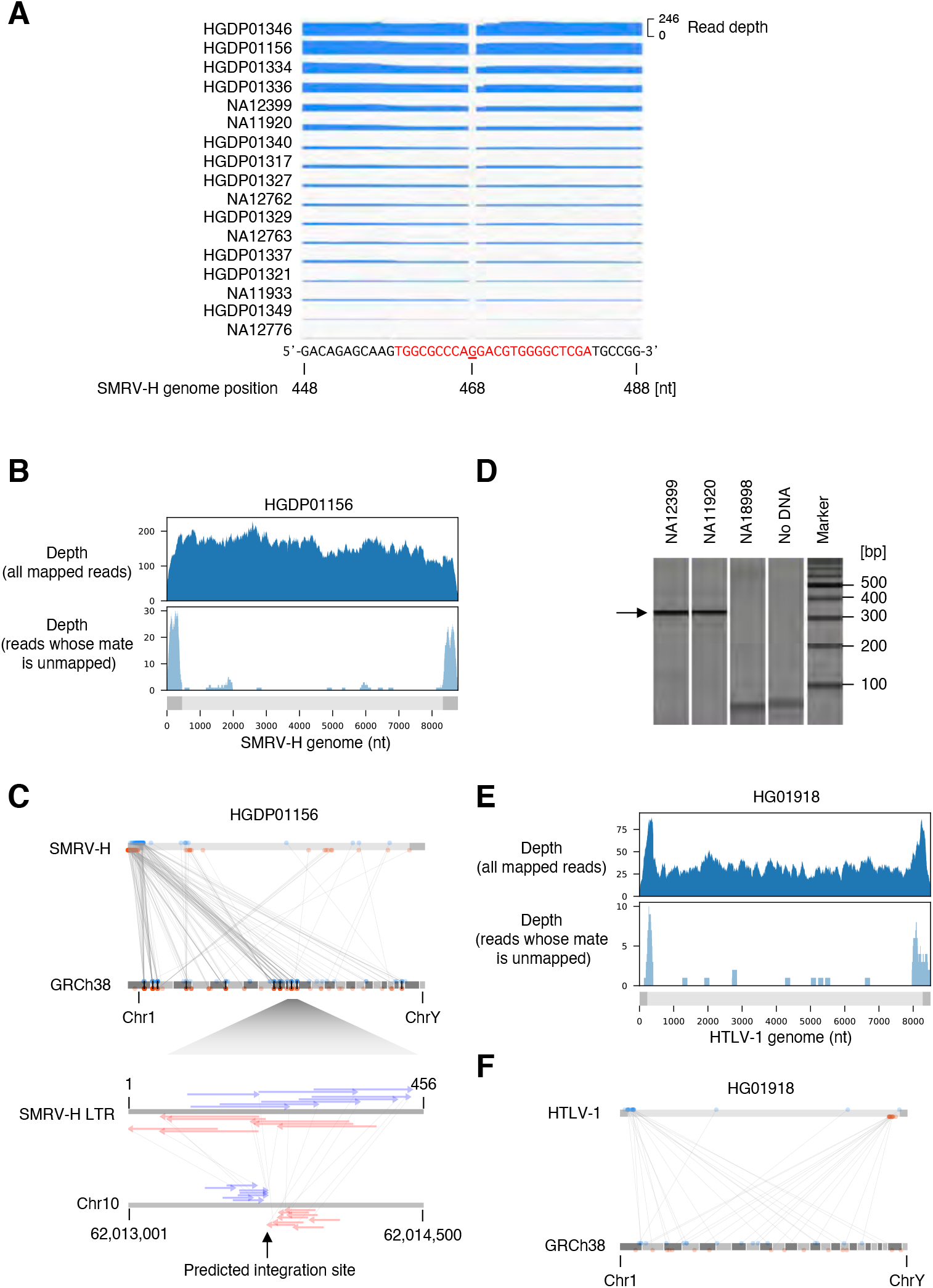
Chromosomal integrations of SMRV and HTLV-1 A. Depth of WGS reads mapping to the primer binding site (PBS) of the SMRV-H genome. Seventeen datasets with at least one read mapping to PBS are shown. One dataset did not have any read mapping to PBS. The PBS of SMRV-H is shown with red characters. In all WGS datasets, the SMRV reads lack the guanosine present at the 468th nucleotide of the SMRV-H genome. B. Depth of HGDP01156 reads mapping to SMRV-H. Upper panel shows the depth of all reads in the dataset mapping to the SMRV-H genome. Lower panel shows the depth of reads mapping to the SMRV-H genome whose mate is not mapped to the SMRV-H genome. Virus genome structure is shown as gray bars. LTR are shown as dark gray rectangles. C. Mapping positions of SMRV-chromosome hybrid reads. Read-1 and Read-2 of a read-pair are connected with a line. All LTR-mapped reads are shown on the left LTR. The lower panel shows the predicted SMRV integration site on chromosome 10. Gray bar in the top of the upper panel represents the virus genome structure. Dark gray rectangles represent LTR. Reads mapping to the forward and reverse directions are shown as blue and red arrows, respectively. D. Detection of SMRV DNA from 1kGP LCLs by PCR. Genomic DNA extracted from the indicated LCLs were used as templates for PCR. WGS datasets from NA12399 and NA11920 are positive for SMRV, while that of NA18998 is negative. E. Depth of HG01918 reads mapping to HTLV-1. Upper panel shows the depth of all reads in the dataset mapping to HTLV-1. Lower panel shows the depth of reads whose pair is not mapped to the HTLV-1 genome. F. Mapping positions of HTLV-1-chromosome hybrid reads. Read-1 and Read-2 of a read-pair are connected with a line. The reads mapping to left LTR was kept when a read was multi-mapped to both left and right LTR. The genome position of reads mapping only to right LTR were replaced to the left LTR.

### Human immunodeficiency virus 1 and human T-lymphotropic virus 1

We detected reads mapping to HIV-1, whose primary targets in the peripheral blood are T lineage cells, in 8 datasets with a maximum coverage of the viral genome 8.5% and depth 0.29x, inconsistent with germline integration (Supplementary Figure 2). LCLs are generated by infecting PBMCs with EBV, which infects mature B cells. Accordingly, most LCL B cell receptors (BCR) have undergone V(D)J recombination, the signature of mature B cells. Moreover, the mode of BCR clonality in a subset of the LCLs analyzed here is one; i.e. they are monoclonal (29). Expression of a rearranged T cell receptor, consistent with presence of T lineage cells, was observed in only one LCL among over 450 screened. HIV-1-mapped reads thus likely either result from infection of hematopoietic progenitor cells (30), ongoing infection of LCLs (31), or from contamination. We did not attempt to confirm the presence of infectious HIV-1 from these cell lines.

Like HIV-1, HTLV-1 is a known human pathogen endemic in populations studied here. HTLV-1 is often transmitted perinatally; analyzing WGS is thus an opportunity to distinguish somatic vertical transmission from potential occult germline horizontal transfer of HTLV-1. Five datasets showed HTLV-1 reads, with read depths ranging from 0.03x (a single paired-end read) to 1.1x relative to autosomes. Two datasets contained HTLV-1-mapping reads at a depth potentially consistent with heritable integrations, 0.55x and 1.1x respectively. Using the hybrid reads approach described above, we demonstrated multiple integrations (Figure 2E, F, Supplementary Figure 4), arguing against germline-inherited integration as the cause of the high abundance of HTLV-1-mapped reads in these datasets. Like HIV-1, HTLV-1 is not well known to infect and integrate into B cells, the source of most LCLs. Thus integration of HTLV-1 into hematopoietic progenitor cells and maintenance of integration site diversity through the LCL generation process, or ongoing replication of HTLV-1 in LCLs, may explain these findings. Subjects whose LCLs sequencing datasets suggest presence of SMRV, HIV-1, and HTLV-1 are listed in Supplementary Table 1.

### Human herpesvirus 6

Germline-integrated HHV-6 has been reported in some of the same datasets analyzed here (32), however we recently described another form of integrated HHV-6 in which a single HHV-6 direct repeat (DR) is present (termed “solo-DR”). The solo-DR form presumably reflects recombination between the two DR regions present in the full-length integrated HHV-6 genome leading to excision of the unique portion of the viral genome (12). Abundant HHV-6 reads were present in 18 datasets (Table 1, Figure 3A, Supplementary Figure 6), suggesting that these subjects likely have chromosomally-integrated copies of HHV-6. One of these samples contained reads mapped only to the DR region of HHV-6B, characteristic of the solo-DR form of integrated HHV-6.

**Table 1.** Summary of integrated HHV-6 identified from 1kGP and HGDP

| Database | Sample | HHV-6 | Structure | Population | Reference |
| --- | --- | --- | --- | --- | --- |
| 1kGP phase3 | HG00245 | HHV-6B | Full | GBR/EUR | Telford *et al.* |
| 1kGP phase3 | HG00362 | HHV-6B | Full | FIN/EUR | Telford *et al.* |
| 1kGP phase3 | HG01058 | HHV-6B | Full | PUR/AMR | Telford *et al.* |
| 1kGP phase3 | HG01162 | HHV-6B | Full | PUR/AMR | Telford *et al.* |
| 1kGP phase3 | HG02016 | HHV-6B | Full | KHV/EAS | Telford *et al.* |
| 1kGP phase3 | HG02301 | HHV-6B | Full | PEL/AMR | Telford *et al.* |
| 1kGP phase3 | HG00145 | HHV-6B | Solo-DR | GBR/EUR | This study |
| 1kGP phase3 | HG00657 | HHV-6A | Full | CHS/EAS | Telford *et al.* |
| 1kGP phase3 | HG01277 | HHV-6A | Full | CLM/AMR | Telford *et al.* |
| 1kGP phase3 | NA18999 | HHV-6A | Full | JPT/EAS | Zhang *et al.* |
| 1kGP pilot | NA19381 | HHV-6B | Full | LWK/AFR | Telford *et al.* |
| 1kGP pilot | NA19382 | HHV-6B | Full | LWK/AFR | Telford *et al.* |
| HGDP | HGDP00092 | HHV-6B | Full | Balochi/SAS | Zhang *et al.* |
| HGDP | HGDP00614 | HHV-6A | Full | Bedouin/MES | This study |
| HGDP | HGDP00628 | HHV-6A | Full | Bedouin/MES | This study |
| HGDP | HGDP00800 | HHV-6B | Full | Orcadian/EUR | This study |
| HGDP | HGDP00802 | HHV-6B | Full | Orcadian/EUR | This study |
| HGDP | HGDP00813 | HHV-6B | Full | Han/EAS | Zhang *et al.* |
| HGDP | HGDP01065 | HHV-6B | Full | Sardinian/EUR | Zhang *et al.* |
| HGDP | HGDP01077 | HHV-6B | Full | Sardinian/EUR | Zhang *et al.* |

**Figure 3.**
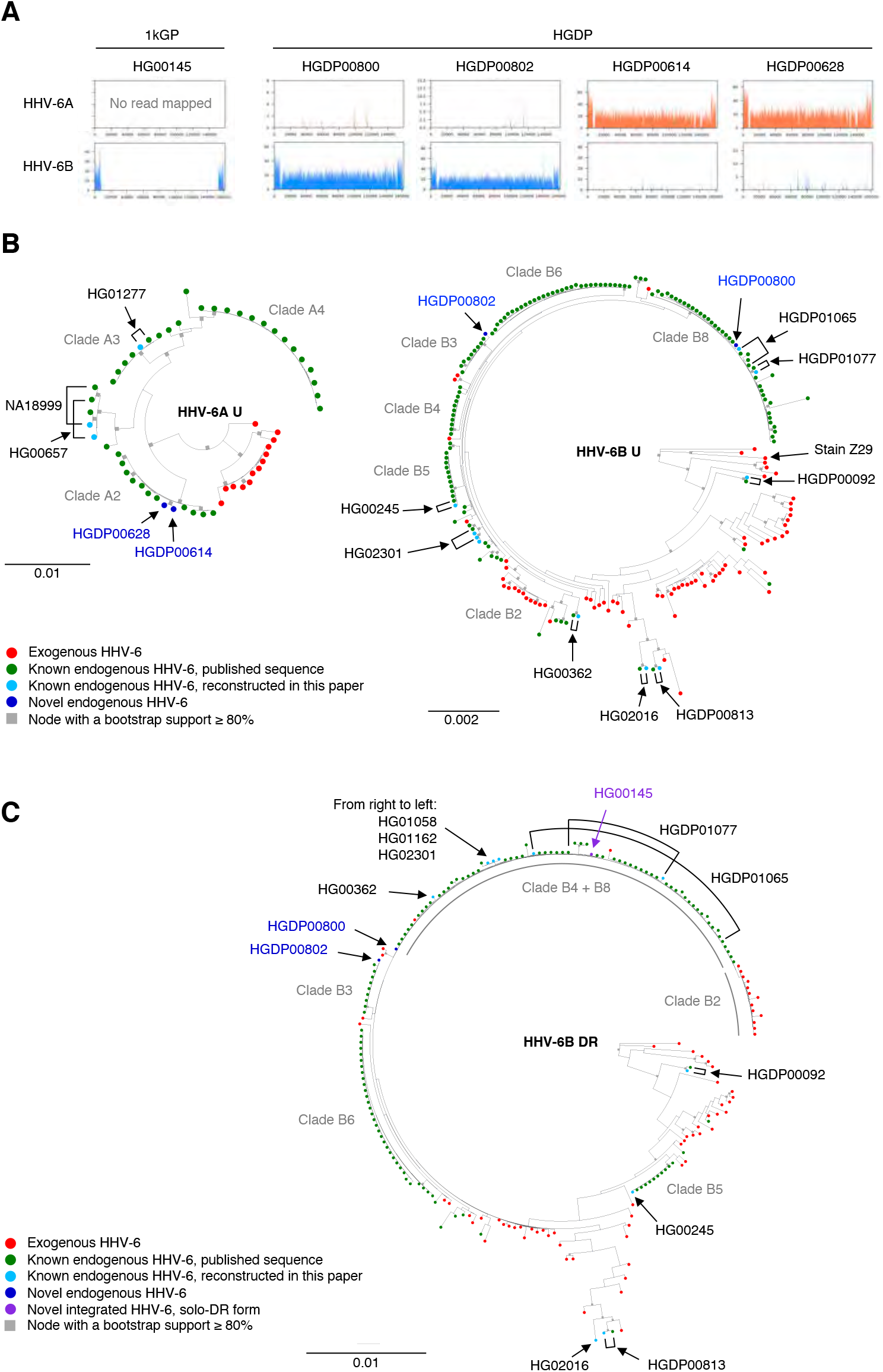
Detection and phylogenetic analysis of endogenous HHV-6. A. Depth of reads mapping to HHV-6 in the 5 WGS datasets from 1kGP and HGDP. B. Phylogenetic trees inferred from U regions of HHV-6A and B. The publicly available sequences of endogenous and exogenous HHV-6 as well as ones reconstructed in the present study were used. C. Phylogenetic tree inferred from DR regions of HHV-6B. The publicly available sequences of endogenous and exogenous HHV-6B, as well as ones reconstructed in the present study, were used. B, C. Clade names defined in the phylogenetic analysis in Aswad *et al*. are shown.

To understand the origin of these integrated HHV-6 variants, we analyzed their relationship to previously-reported exogenous and integrated HHV-6 sequences. All reconstructed sequences clustered with previously-reported sequences (Figure 3B, Supplementary Figure 8, 9). Two endogenous HHV-6A found in Bedouin subjects clustered with clade A2 sequences, which were previously found in subjects in the US and UK, suggesting these subjects share endogenous HHV-6A derived from a single integration event. Shared ancestry of this chromosomal fragment, previously shown to correspond to the telomere of chromosome 17p, between subjects on three continents is consistent with the deep evolutionary relationship of endogenous HHV-6A (13). Newly-reported endogenous HHV-6B in two subjects in HGDP were grouped with clade B8 and Clade B3, respectively. Clade B8 integrations also map to chromosome 17p, which bears a short telomere (33). The integration site of clade B3 endogenous HHV-6 has not yet been determined.

To clarify the origin of the newly-detected solo-DR variant, we generated a phylogenetic tree using only DR sequences (Figure 3C, Supplementary Figure 10). The solo-DR form reconstructed from the HG00145 genome was present in the same clade as DRs from clade B4 and B8 full-length endogenous HHV-6B. This suggests that the solo-DR form likely arose from an HHV-6B source closely related to that of clade B4 and B8, but precludes confident inference that the solo-DR represents a germline rearrangement of a full-length endogenous HHV-6. Detection of solo-DR integrated HHV-6 in this already well-characterized dataset shows that screening WGS databases may provide additional information regarding the excision and potential for reactivation of endogenous HHV-6.

### Human endogenous retrovirus K

Previous studies of the DNA virome detectable from human genome sequencing datasets have noted inter-individual variation in reads mapped to HERV-K (18, 19). This variation (e.g. Figure 1) has been speculated to result from polymorphic HERV-K integrations (19). While the viruses described above are absent from reference human genomes, HERV-K is present in multiple nearly-identical copies in reference genomes. This makes detecting additional non-reference integrations and mapping the chromosomal location of polymorphisms challenging. Previous advances in mapping HERV-K polymorphisms have been made by local breakpoint reconstruction using read-pairs that span insertion junctions (2). Taking a different approach to this problem, we extracted all unique *k*-mers of length 50 from the aligned portion of reads mapped to the HERV-K113 provirus (i.e. excluding the LTRs, Figure 4A). To filter only those *k*-mers derived from HERV-K loci that are polymorphic between humans, we extracted *k*-mers which were absent in at least one subject and present in at least two subjects (n=37,845 “presence-absence type” *k*-mers). Hierarchical clustering of presence/absence-type *k*-mer occurrences recapitulates the continental human population supergroups (Figure 4B), as does clustering based on the allele frequency of previously reported polymorphic HERV-K in human subpopulations (2). This suggested that presence-absence *k*-mers may be a suitable proxy to allow for discovery of additional polymorphic HERV-K alleles.

**Figure 4.**
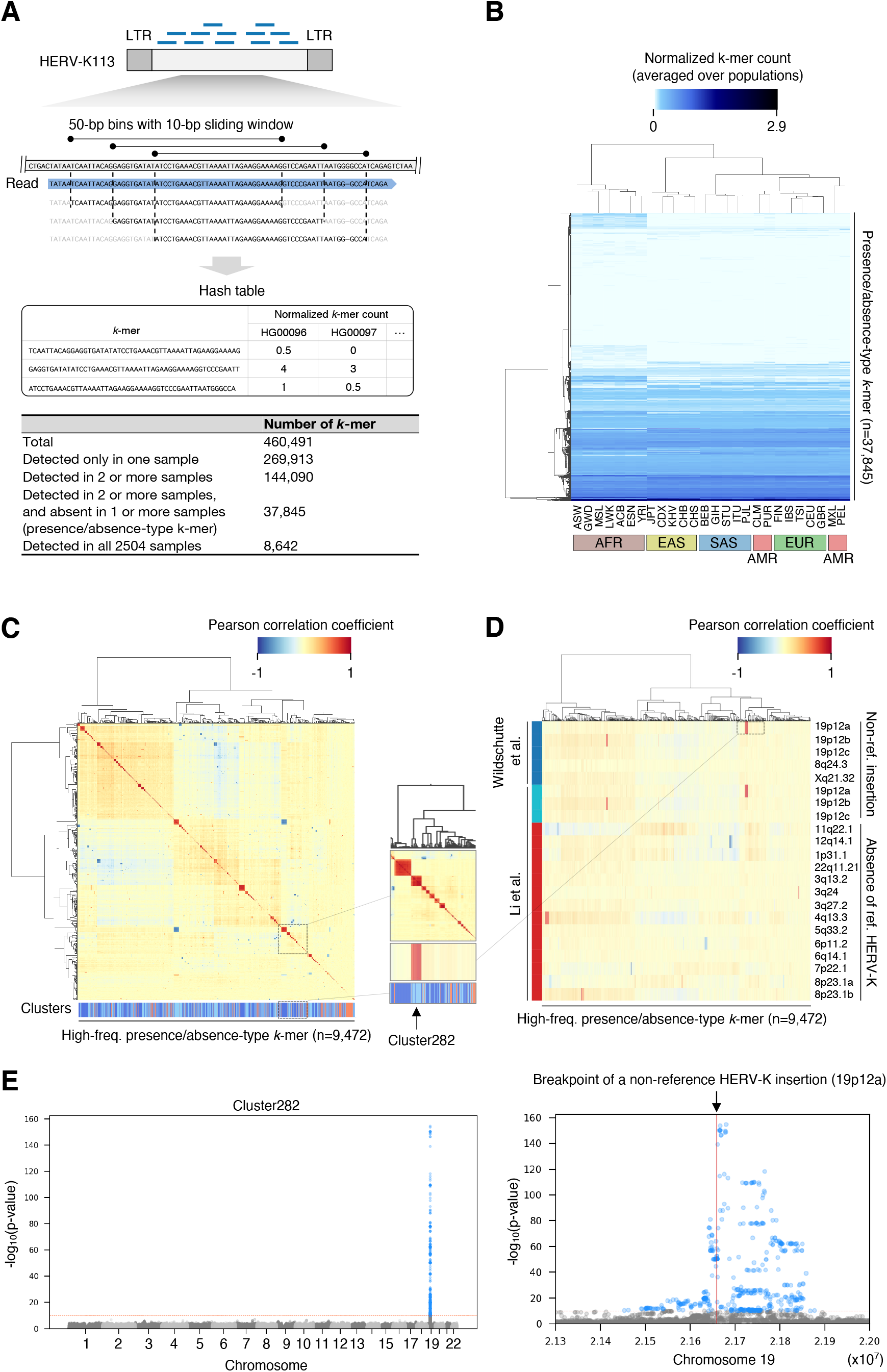
HERV-K *k*-mer detection from 1,000 Genomes Project WGS A. Schematic representation of *k*-mer counting from WGS reads mapping to HERV-K113. The HERV-K113 genome was split into 50-bp bins with a 10-bp sliding window, then, sequences of the mapped reads corresponding to the HERV-K 50-bp bins were listed. The lower table shows the number of *k*-mers detected from 2,504 WGS datasets from the 1,000 Genomes Project. B. Hierarchical clustering of *k*-mers based on their frequencies in 26 populations. Heatmap shows the normalized *k*-mer count averaged over populations. C. Clustering of presence-absence type *k*-mers by Pearson correlation coefficient. Clustering were performed by Ward’s method (upper heatmap) and DBSCAN (lower color-bar). The heatmap shows the Pearson correlation coefficient between *k*-mers, and the lower color-bar shows the clusters. Neighboring *k*-mer clusters are shown as either dark or light blue. Orange represents the *k*-mers which were not clustered. D. Correlation between the occurrence of HERV-K *k*-mers and previously reported HERV-K polymorphisms. Heatmap shows the Pearson correlation coefficient between the presence of *k*-mers and polymorphic HERV-K reported in the two previous studies. (C, D) Insets in the panel C and D shows that the occurrence of *k*-mers in cluster282 have high correlation to the presence of known polymorphic HERV-K in 19p12a. E. GWA using occurrence of *k*-mers detects known polymorphic HERV-K. Manhattan plots show SNVs with association to the occurrence of *k*-mers in the cluster282. SNVs with p-value lower than 8.33e-11 are shown as blue dots. Red solid line in the right panel shows the position of known non-reference HERV-K.

We hypothesized that structurally-polymorphic HERV-K alleles generate multiple unique *k*-mers with the same pattern of presence or absence in multiple subjects. To test this, we generated an all-by-all Pearson correlation coefficient matrix for presence/absence type *k*-mers which were detected in more than 50 subjects (n=8,642) then performed hierarchical clustering of *k*-mers. This revealed multiple groups of *k*-mers with presence/absence patterns that were highly correlated with one other (Figure 4C), suggesting that a single polymorphic HERV-K locus could generate multiple presence/absence-type *k*-mers. We formally defined clusters using DBSCAN (see methods), resulting in 597 clusters of highly co-associated *k*-mers.

We next investigated how the observed clusters of presence/absence-type *k*-mers relate to known HERV-K polymorphisms. Some *k*-mer clusters correspond very well with those of known HERV-K polymorphisms previously described in these same subjects (Figure 4D). For example, the presence/absence pattern of *k*-mers in cluster282 is highly correlated with that of a non-reference HERV-K insertion on chr19. This suggests that in some cases, clusters of presence/absence-type *k*-mers reflect HERV-K polymorphisms.

Determining where on the chromosome a particular polymorphic repetitive genetic element is located can be challenging using short read sequencing data, because the read evincing a polymorphism could potentially have arisen from a number of loci bearing nearly identical elements. In addition, while some reads mapping to HERV-K LTRs are paired with uniquely-mappable reads from flanking non-repeat DNA, variant reads mapping to the HERV-K provirus are rarely paired with uniquely-mappable non-repetitive reads, as a consequence of the size of most sequencing library inserts. To overcome these challenges, we took advantage of the linkage disequilibrium (LD) structure of human chromosomes. We hypothesized that a *bona fide* polymorphic HERV-K element, giving rise to a *k*-mer cluster, would be in linkage disequilibrium with nearby SNVs. If so, analyzing genome-wide association (GWA) with SNVs would allow us to locate the polymorphic HERV-K. To validate whether this approach is able to accurately report the genome positions of polymorphic HERV-K, we examined a known non-reference HERV-K insertion. GWA analysis using the presence/absence-pattern of cluster282 *k*-mers, considered as a binary trait, detected a significant association with a single approximately 300-kb region on chromosome 19 known to contain the non-reference HERV-K insertion (Figure 4E). This approach of using <u>l</u>inkage <u>d</u>isequilibrium to <u>f</u>ind <u>r</u>epetitive <u>e</u>lement <u>d</u>ifferences, an approach which may be applicable to other repetitive elements, is abbreviated here as “LDfred.”

We performed GWA analyses using the presence/absence patterns of all 597 clusters. As a result, 503 clusters detected at least one genome region with a Bonferroni-corrected genome-wide significant association; clusters with associated regions tended to consist of more *k*-mers and/or be present in more subjects (Supplementary Figure 11). We merged clusters that were associated with overlapping regions (see methods), resulting in a total of 79 HERV-K *k*-mer-associated loci spanning a total of 74.7 Mb. These loci most often include regions in which the *P* values peak sharply, pinpointing the most tightly-linked LD block and narrowing the presumed location of the HERV-K variant (e.g. Figure 4E). Consistent with previous work showing that mobile elements are often linked to trait-associated SNVs (34, 35), the SNVs comprising these HERV-K polymorphism-associated haplotypes are associated with numerous human traits (Supplementaly Table 2), including 5 loci in which SNVs from the GWAS catalog (36) overlap precisely with the genomic regions evincing polymorphic HERV-K (Supplemental Figure 12). To check whether these new *k*-mer-associated loci indeed contain polymorphic HERV-K, we evaluated overlap with known polymorphic HERV-K loci (2, 4–6). Seven out of 10 known non-reference proviruses are present within the observed loci (2 known non-reference proviruses are not on the reference autosomes included for GWA), as are all 16 reference HERV-K known to be absent in some subjects (Figure 5).

**Figure 5.**
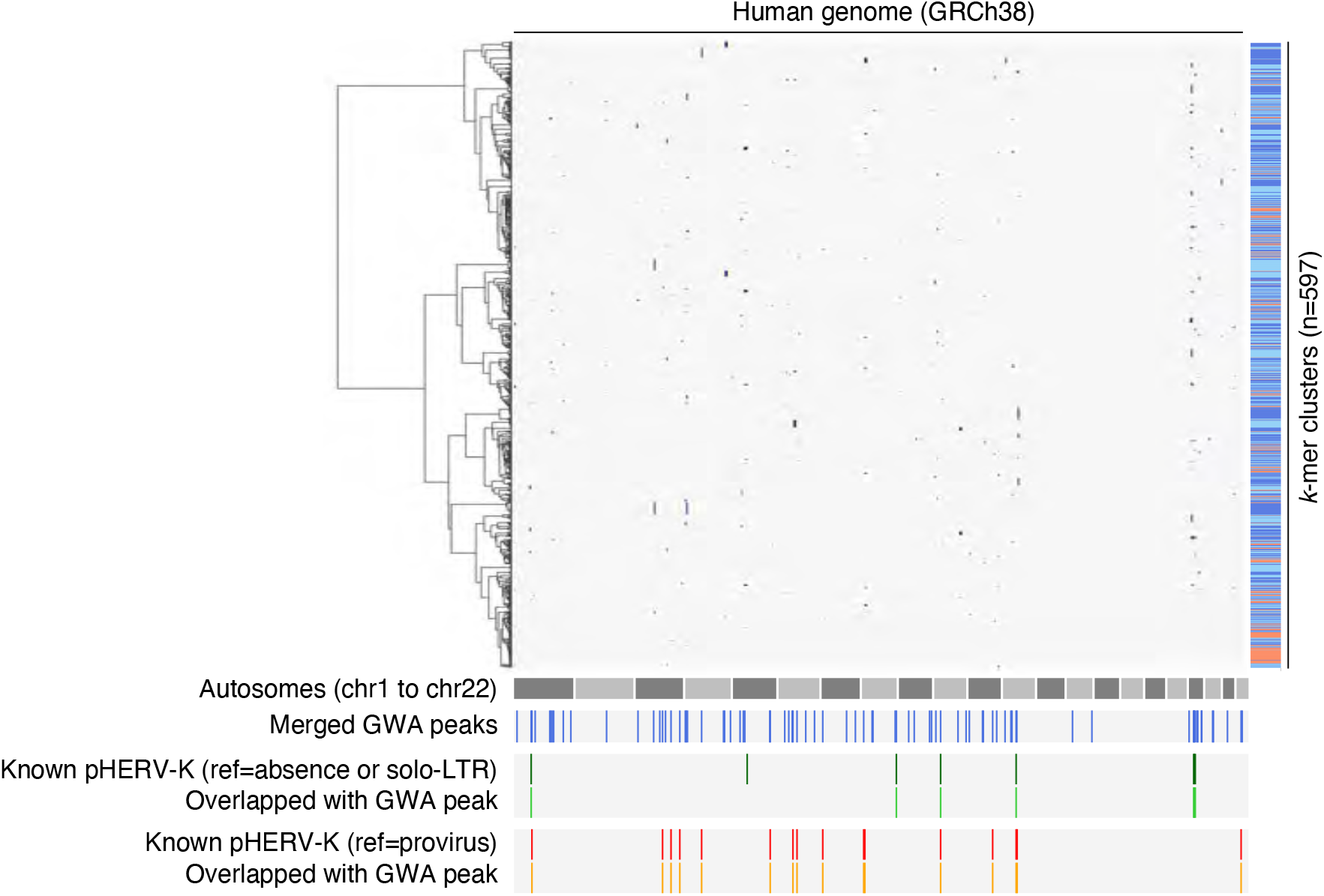
Genome positions associated with HERV-K *k*-mers The blue dots in the left clustermap show the genome positions with association to *k*-mer clusters by GWA analysis. The blue lines in the third right column show the 79 genome loci associated with *k*-mer clusters. The dark green lines show the 8 known non-reference HERV-K on autosomes. The light green lines show the 7 *k*-mer cluster-associating genome loci overlapping with the known non-reference polymorphic HERV-K on autosomes. The dark orange lines show the 16 known reference polymorphic HERV-K on autosomes. The light orange lines show the 16 *k*-mer cluster-associating genome loci overlapping with the known reference absent HERV-K on autosomes.

To assess whether LDfred could localize previously unknown polymorphisms, we checked long-read sequencing datasets for reference HERV-K that are absent in some subjects. We filtered an extensive catalog of SVs in three subjects, generated using multiple sequencing technologies, for deletions that overlap with HERV-K (37). We found 24 reference HERV-K elements with evidence of full or partial absence in at least one of the three subjects (Supplementary Table 3). These 24 SVs spanned from 72 to 9,468-bp. We checked if these HERV-K SVs were present in loci associated with *k*-mer clusters, and whether the presence/absence pattern of the *k*-mers in each cluster was consistent with the presence/absence of SV in the three subjects (see methods). Of the 24 HERV-K SVs, 9 were concordantly detected (Supplementary Table 4); concordantly detected SVs tended to be longer than those not concordantly detected. Notably, 4 out of 9 detected HERV-K SVs have not been previously reported as polymorphic HERV-K. These 4 unreported SVs are attributable to recombination between LTRs (Supplementary Figure 13), which is particularly difficult to find using existing algorithms and short-read sequencing. This demonstrated that LDfred can localize unknown HERV-K provirus polymorphisms, including provirus/solo-LTR polymorphisms.

SVs in complex or duplicated genome regions are also difficult to identify using short-read data and available methods (38). To check the utility of our approach for this purpose, we focused on HERV-K loci at chromosome ends, known to be complex genome regions (39). One locus, associated with cluster352, is in the subtelomeric region of the short arm of chromosome 4. There are no HERV-K proviruses in this region, however there are 2 solo-LTRs, suggesting the possibility of a provirus/solo-LTR polymorphism, or an additional non-reference HERV-K insertion. We assessed whether either of these reference LTR loci sometimes contain a provirus using long read sequencing data from a trio (40). WGS from the father and child contained cluster352 *k*-mers, but the mother did not harbor any *k*-mers in cluster 352, suggesting that the father and child could carry a non-reference provirus. We inspected reads mapped to the subtelomere of chromosome 4 and found a read containing non-reference provirus at the region corresponding to one of the solo-LTRs in the reference genome at this locus (Figure 6). Thus LDfred can detect provirus/solo-LTR polymorphic HERV-K loci in complex regions including subtelomeres.

**Figure 6.**
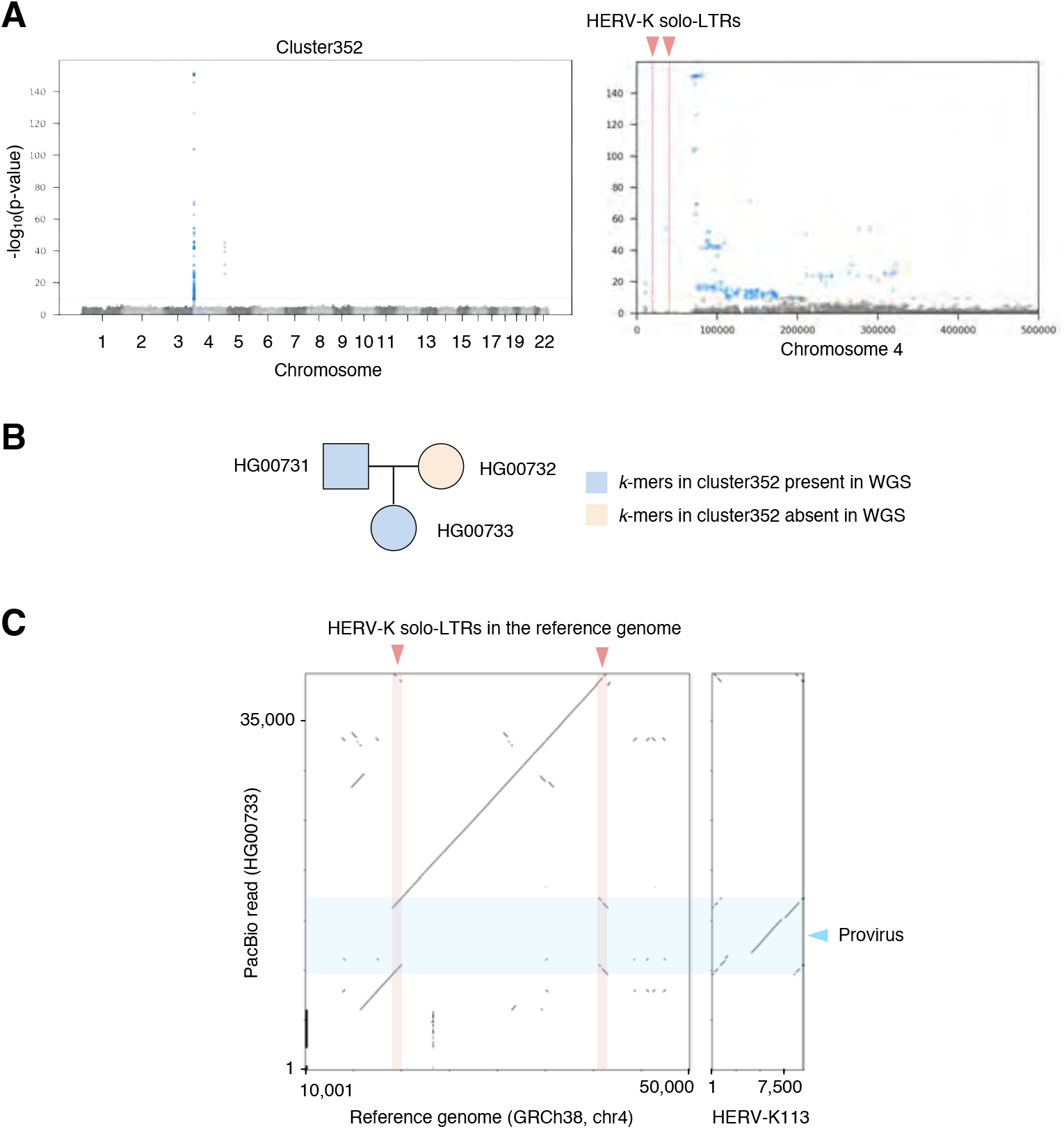
*K*-mer-based method detects previously-unreported HERV-K polymorphism in subtelomere A. SNVs associating with *k*-mers in cluster352. The right panel shows the region near the end of the p-arm of the chromosome 4. SNVs with p-value lower than 8.33e-11 are shown as blue dots. The red solid lines in the right panel show two reference HERV-K solo-LTR in the subtelomere region. B. Presence and absence of *k*-mers of cluster352 in the public high-coverage short-read WGS of the Chinese trio. C. HG00733 contains non-reference provirus. Left panel shows the dot matrix between the reference human genome and a PacBio read of HG00733. The right panel shows the dot matrix between the reference HERV-K113 and the PacBio reads of HG00733. Light blue and light red rectangles represent HERV-K provirus and solo-LTR, respectively.

Of the 79 loci identified here, five loci contain known non-reference presence/absence or provirus/solo-LTR polymorphisms and 14 contain reference HERV-K known to be absent in some subjects (including some merged loci containing more than one polymorphism). Six loci reflect provirus/solo-LTR polymorphisms that have not been previously reported, but which are readily demonstrated in long-read data (Figure 6, Supplementary Figure 13, 14). The remaining loci cannot be assessed using available long-read data, because the minor variants, as determined by *k*-mer pattern, are not present in subjects with available data. To determine the polymorphisms giving rise to the signal that allowed us to identify the remaining loci, we chose three for which the regions flanking reference HERV-K in these regions allowed us to design specific primers. Targeted long-read sequencing revealed differences in the HERV-K at these loci consistent with the *k*-mer pattern differences between the individuals. The nature of the variation at these loci was not structural; instead, it consisted of multiple SNVs (up to 11) linked in a haplotype (Figure 7; Supplementary Figure 15, 16). This high degree of linked SNV variation distinguishing HERV-K proviruses at the same locus is unexpected due to accrual of substitutions; 11 mutations across these 4.9 kilobases of the HERV-K provirus (Figure 7) would accumulate over 4.4 million years (41), and the linkage between them would be expected to be degraded by crossover recombination events during that period. Instead, this more likely reflects interlocus gene conversion via recombination, which has previously been described for HERV on the basis of comparison of LTRs in different species (42), or introgression. Notably, the specific HERV-K haplotypes present as minor variants in these loci are not detected by BLASTn search of the hg38 reference genome. Thus LDfred can localize previously-uncharacterized sources of non-structural HERV-K variation.

**Figure 7.**
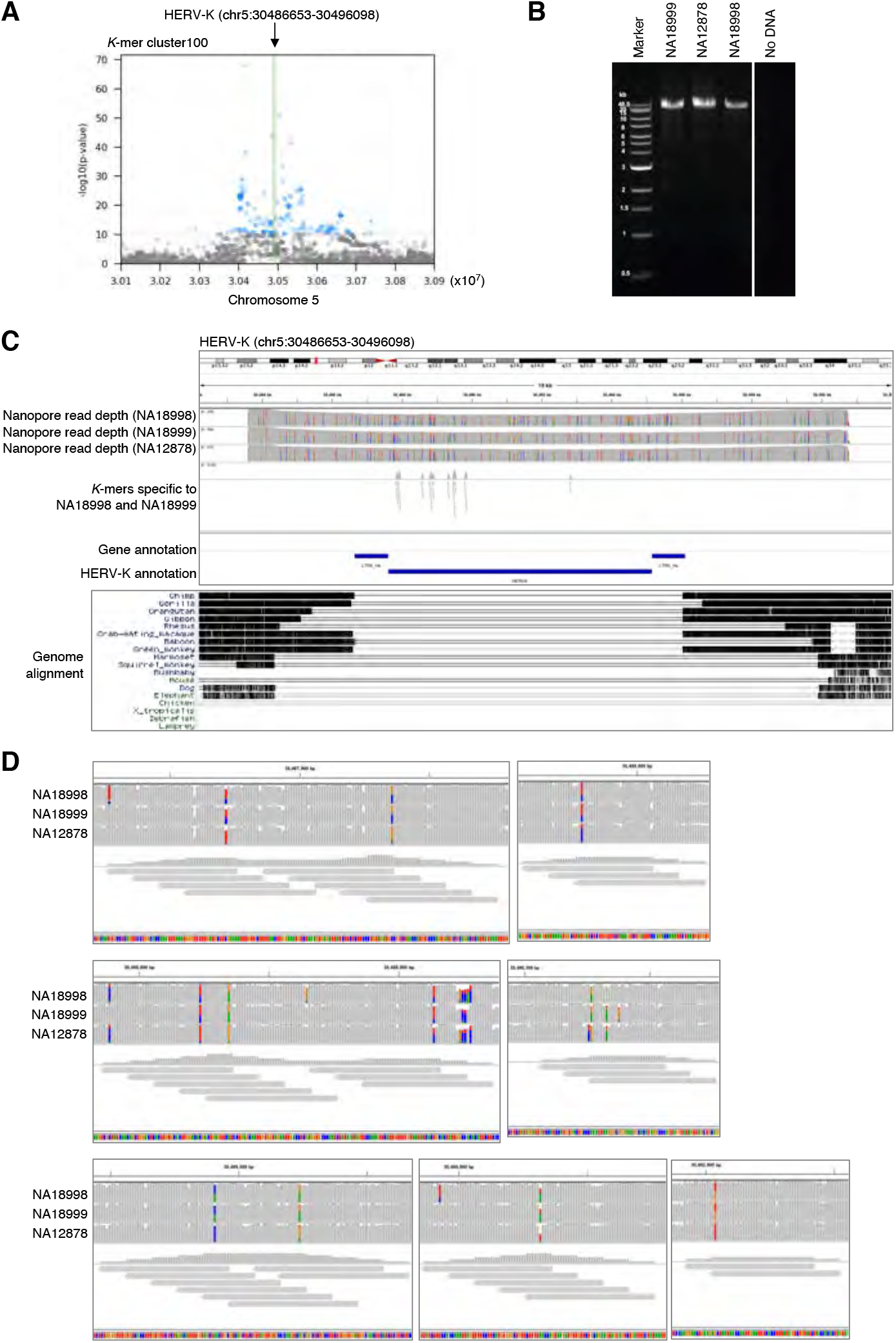
Potential interlocus gene conversion in HERV-K localized by LDfred A. Manhattan plot showing SNVs associating the *k*-mer cluster100. SNVs with p-value lower than 8.33e-11 are shown as blue dots. Green line shows the reference HERV-K provirus. B. Amplification of the HERV-K provirus by PCR. HERV-K provirus with adjacent sequence was amplified and PCR products were separated by gel electrophoresis. DNA extracted from LCLs originating from NA18998, NA18999, and NA12878 were used as templates. C. Upper panel: IGV view of long-read sequencing reads mapping to HERV-K. The PCR amplicons were sequenced using an Oxford Nanopore flongle flow cell and mapped to GRCh38. *k*-mers in *k*-mer detecting the HERV-K were also mapped to the PCR target regions. Lower panel: UCSC genome browser view showing the Multiz Alignment of 100 Vertebrates track. D. Enlarged images of panel C. NA12878 carries two alleles of a non-reference HERV-K haplotype (which is not observed elsewhere in the reference genome) also present as a single allele in NA18998 and NA18999.

## Discussion

This work provides a comprehensive picture of virus-derived structural variations in two well-studied global WGS datasets. We found previously-missed germline structural variants arising from HHV-6 and HERV-K, as well as virus integration in somatic cells due to natural infection or contamination. The presence of SMRV integrations in LCLs introduces caveats in analyzing these materials and the data derived from them. This is especially notable for cells used in the 1kGP, from which only subjects from Utah (collected by CEPH) are SMRV-positive. In most subjects, the number of virus-chromosome hybrid reads detected at each specific genome locus is low, suggesting a mosaic of cells with different integration events. However, in some datasets, such as HGDP01156 and HGDP01346, a substantial number of hybrid reads are detected from individual genome loci, suggesting a high fraction of clonal cells bearing the same virus integration event. In such cases, the SMRV insertion could influence nearby variant calls, and could also influence clonal expansion during the course of LCL culture (43). SMRV is transcribed in SMRV-positive LCLs, and this viral RNA could influence host transcription, for example by triggering innate immune pattern recognition receptors. However, we observed no significant change in expression of interferon-stimulated genes (ISGs). This may be related to the concurrent presence of EBV, reported to counteract ISGs, in these cells (44). SMRV-positive LCLs did express a few genes significantly differently than SMRV-negative cells, which has implications for interpreting the results of studies using these cells and datasets (28). We confirmed the presence of SMRV DNA in recently-distributed LCLs from these donors. While biosafety regulations vary by locality, our results reinforce that even well-characterized LCLs should be handled as potential sources of infectious viruses.

We found evidence of infection of LCL progenitors with HIV-1 or HLTV-1. This was unexpected because B cells, the proximal progenitor of most LCLs, are not efficiently infected by either of these viruses. However EBV transformation has recently been shown to permit replication of some types of HIV-1 in B cells (31). Given the diversity of integration sites observed, we suspect that a similar phenomenon may explain the presence of HTLV-1 in LCLs, although we cannot exclude somatic mosaicism due to infection of hematopoetic stem cells (45). We found no evidence of germline structural variants related to HIV-1 or HTLV-1. As has recently been noted, HIV-1 is capable of infecting germ cells (46). Our result should be interpreted to indicate that the plausible upper bounds of the global allele frequency of such variants is ~0.01%; larger-scale projects, or projects sequencing populations with higher prevalence of infection by these viruses, could apply the methods used here to discover these rare variants, if they exist.

An endogenous form of Koala retrovirus has been considered a unique opportunity to study a virus that is in the process of endogenization in mammals (47). Our report clarifies the extent to which humans also harbor a virus that straddles the endogenous/exogenous divide, HHV-6. Understanding the relationship between endogenous and exogenous HHV-6 is a critical question to which the tools presented here can be usefully applied. The solo-DR form of integrated HHV-6B is present in one 1kGP dataset, but was not detected in previous surveys of these data and samples (32). The presence of unreported solo-DR integrated HHV-6 in such a well-investigated database suggests that the prevalence and diversity of integrated HHV-6 has likely been underestimated in other studies as well. Phylogenetic analysis suggests that this solo-DR variant was potentially formed by partial excision of B4 or B8 clade endogenous HHV-6B by recombination, leaving behind one DR as a “scar.” This molecular event has been proposed to lead to viral reactivation. In that context, it is notable that a few reportedly-exogenous HHV-6B sequences are quite closely related to B4 and B8 endogenous HHV-6B, while the majority of sampled exogenous HHV-6B are more divergent. We cannot exclude the possibility that this solo-DR variant is related to a third independent integration by a virus related to those giving rise to B4 and B8 endogenous HHV-6B. Furthermore, while the DR was evidently present in a hemizygous state in the nuclei of LCLs from this subject, there is no evidence that it is endogenous or “inherited” as is often used to describe such variants; we consider it most accurate to describe it as chromosomally-integrated HHV-6 until another identical-by-descent variant should happen to be observed, integrated in another human’s genome.

We recently reported the presence of the solo-DR form of endogenous HHV-6 in the Japanese population. The solo-DR positive subject in 1kGP is of European ancestry, demonstrating that solo-DR variants are present in populations of different ancestry. Integrated HHV-6 has been associated with angina pectoris and pre-eclampsia, and in both cases a virus-dependent mechanism has been postulated (48, 49). It is thus important to ask whether the solo-DR form of endogenous HHV-6, or only full length HHV-6 integration, is associated with human phenotypes and diseases. Screening additional databases using the tools developed here can capture the complete diversity of HHV-6 integration in human chromosomes, leading to a deeper understanding of its potential influence on human diseases. We also found four previously unreported full-length endogenous HHV-6. Among these are independent integrations, one HHV-6A and one HHV-6B, into chromosome 17p, which is reported to carry the shortest telomere of the 46 chromosome arms (33). This mirrors our observation in the Japanese population, in which two prevalent endogenous HHV-6 variants, one HHV-6A and one HHV-6B, are integrated on chromosome 22q, which carries the second shortest telomere, on average (33). One full-length endogenous HHV-6B falls within Clade B3, the integration site of which is currently unknown, but will be able to be mapped using this LCL in the future.

We also explored the polymorphisms of HERV-K, which includes the most recently integrated HERVs. The overlap of GWA peaks with genome loci known to harbor polymorphic HERV-K suggests that our approach captures HERV-K polymorphisms that can be discovered by other methods. In addition, our approach identifies regions with association to HERV-K *k*-mers that were not previously reported to be polymorphic; these loci likely contain unreported HERV-K polymorphisms. Using long-read sequencing data, we confirmed that six of these loci indeed contain HERV-K structural variants. The *k*-mer-based method presented here does not explicitly distinguish presence/absence-type polymorphisms from SNV polymorphisms in the non-LTR portion of HERV-K. *K*-mer signals due to individual substitutions, as would begin to accumulate after integration of a full length HERV-K provirus at a given locus, may also be detected. This approach conceptually allows finding any sequence differences between repeat element loci, and may be useful for other difficult-to-map repeat elements (38, 50). In total we document the nature of the HERV-K polymorphisms explaining over 20 of the loci reported here, yet many loci remain to be validated; increasing use of long-read sequencing should enable this soon (51).

We set a threshold of clustering only presence/absence type *k*-mers found in at least 50 subjects, so HERV-K polymorphisms with allele frequencies below 0.02 should not be detected. While we set this threshold arbitrarily and conservatively, this approach is limited in its ability to localize very rare polymorphisms due to the nature of linkage disequilibrium and the decreased statistical power of association testing using few polymorphism-bearing subjects. We treated *k*-mer presence-absence pattern as a binary trait, yet retaining the continuous variation in these patterns could approximate genotyping, potentially improving localization of polymorphisms in repeats in the future. Previous studies reported that the majority of unfixed HERV-K in humans are solo-LTR type (2). We defined *k*-mers using only reads mapped to the non-LTR regions of HERV-K due to the high sequence similarity between HERV-K LTR and SVA retrotransposons. This enabled us to detect new solo-LTR vs full provirus polymorphisms, but we could not detect solo-LTR vs empty site polymorphisms. To understand polymorphisms of HERV-K comprehensively, including presence/absence of solo-LTR, we will need to expand the *k*-mer-based method; this will cross-detect SVA retrotransposons which are not virus-derived in this same sense as the other variants considered within the scope of this report.

We used targeted long read sequencing to determine the HERV-K polymorphisms present within several of the newly-identified loci. We observed divergent HERV-K haplotypes, differing by 11 linked SNVs within 4,932 bp from the major allele in the most divergent, present at the same locus. This degree of variation at a syntenic HERV-K integration site, absent in other great ape genomes, is unexpected as a result of clock-like accumulation of mutations (52). Among the potential explanations for this phenomenon, two are plausible and warrant discussion. First, a process of non-allelic homologous recombination, most often referred to as gene conversion in this context, could exchange the HERV-K haplotype at the locus *en bloc* with that from another locus. This has often been invoked to explain differences between HERV loci syntenic between species (42), and has been reported for HERV-K (53). However, the potential “source” haplotype for such a recombination could not be identified in the hg38 reference genome (i.e. by BLASTn search using the variant *k*-mers). We thus cannot distinguish whether the source HERV-K element was itself an insertion that has now been lost or fixed as a solo-LTR, nor can we exclude the possibility of introgression of the HERV-K haplotypes from an archaic source. In any case these results point to a previously unexplored cache of HERV-K diversity in human genomes, and offer a new tool to guide its exploration. Considering that infectious HERV-K sequences can be generated with via recombination between known HERV-K elements (54), these previously hidden HERV-K polymorphisms are particularly relevant to study in relation to human phenotypes. Ongoing and future large-scale population sequencing projects will massively expand the data available to address viral contributions to human genomes, and the tools presented here will enable integration of these analyses into the planned output of these consortia (55).

## Methods

### WGS datasets

High-coverage WGS datasets from 1kGP were downloaded from the following URL: ‘ftp://ftp.1000genomes.ebi.ac.uk/vol1/ftp/data_collections/1000G_2504_high_coverage/’. High-coverage WGS datasets from 1kGP Han Chinese trio were downloaded from the following URL: ‘https://www.internationalgenome.org/data-portal/data-collection/structural-variation’. High-coverage WGS dataset from HGDP were downloaded from the following URL: ‘ftp://ftp.1000genomes.ebi.ac.uk/vol1/ftp/data_collections/HGDP/data/’. The utilization of the high-coverage WGS of multigenerational CEPH/Utah families (phs001872) are authorized by the National Human Genome Research Institute through dbGaP for the following project: “The prevalence, evolution, and health effects of polymorphic endogenous viral elements in human populations.”

### Preparation of reference virus genomes

Reference virus genomes were downloaded from NCBI on April-6-2020. We downloaded three files named ‘ftp://ftp.ncbi.nlm.nih.gov/refseq/release/viral/viral.[1,2,3].1.genomic.fna.gz’. These three files contained 12,182 virus genomes, including phages. These files were concatenated into one file and used as reference virus genomes for further analysis.

### Virus detection and reconstruction from WGS

Reads that did not map to the reference human genome were extracted from WGS BAM or CRAM files using ‵samtools view -f 1 -F 3842 | samtools view -f 12 -F 3328 -‵ command. Then, the unmapped reads were converted to FASTQ format using samtools fastq command. FASTQ reads shorter than 20-nt were removed using a custom Python script and excluded from downstream analysis. Retrieved sequences were then mapped to the reference virus genomes using Hisat2 with ‵--mp 2,1 --no-spliced-alignment‵ options. After mapping, reads estimated to be PCR duplicates were marked using picard MarkDuplicates command. Then, the mapping depth and coverage of each virus was calculated using deeptools bamCoverage command with ‵--binSize 1‵ option. Based on the depth and coverage, we searched for viruses with abundant reads using a custom Python script. We labeled ‘virus_exist’ if 5% of a viral genome was covered at more than 2x read depth. Virus genomes with a ‘virus_exist’ label were then reconstructed by incorporating variations to the reference virus genomes. To reconstruct viruses, variations in the reads mapping to virus genomes were called using gatk HaplotypeCaller. The output vcf files were then normalized using bcftool norm command and the reconstructed virus sequences were generated using bcftools consensus command. Regions without any mapped reads were masked by ‘N’ using a custom Python script. The workflow described here was compiled as a Python pipeline and available from the following GitHub repository: ‘https://github.com/GenomeImmunobiology/Kojima_et_al_2020’.

We detected reads mapping to 634 viruses, including 553 phages, in total (Supplemental figure 1 (the original heat map, showing all viruses by all people)). Phage are ubiquitous in the human virome and thus should not necessarily be excluded as a potential source of horizontal gene flow to humans (56). Phage-mapped reads were often found, however they were present at low depth inconsistent with germline integration into human genomes, and were thus excluded from further analysis. The same was true of most eukaryotic viruses, which showed low average read depth, usually less than 1x across the entire length of the viral genome. This may reflect virus infection in a small proportion of cells, contamination from other samples sequenced on the same machine, or mis-mapping. As the primary goal of this study was to detect potentially heritable virus-derived structural variants, these were not analyzed further.

### Endogenous HHV-6 detection and reconstruction from WGS

We developed a bioinformatic pipeline specialized to detect and reconstruct full-length endogenous HHV-6 as well as solo-DR form, because endogenous HHV-6 has a terminal direct repeat sequence (DR), which is not appropriately reconstructed using the virus detection and reconstruction pipeline described above. We extracted reads that did not map to the human genome and mapped these reads to the reference HHV-6 using the same commands described above. Rather than all viral sequences for HHV-6 reconstruction, we used full-length exogenous HHV-6 genomes NC_001664.4 and NC_000898.1 as HHV-6A and HHV-6B, respectively. We judged whether a WGS dataset contains abundant HHV-6 reads using the same cutoff described above. When abundant HHV-6 reads were detected, we reconstructed the full-length HHV-6 sequence using the same reconstruction protocol as described above. The DR region of a reconstructed full-length genome is not accurate, because reads mapping to DR are mapped to both left DR and right DR. The reads with multimapping have the mapping score 0, being excluded from downstream variant calling. To accurately reconstruct DR, all reads mapping to HHV-6 genomes were re-mapped to DR-only. For this reconstruction, we used nucleotides 1-1089 of NC_001664.4 and 1-8793 of NC_000898.1 as DR-only sequences of HHV-6A and HHV-6B, respectively. The workflow described here was compiled as a Python pipeline and available from the following GitHub repository: ‘https://github.com/GenomeImmunobiology/Kojima_et_al_2020’.

For validation of the accuracy of endogenous HHV-6 reconstruction, we used a 35x coverage WGS dataset from subject NA18999, a Japanese subject known to contain full-length endogenous HHV-6A (Supplementary Figure 7). The reconstructed sequence covered 96% of the reference genome (U1102) with 96.7% similarity. Phylogenetic analysis of the reconstructed sequence with sequences from this subject previously determined by Sanger sequencing demonstrate that the reconstructed sequence is very close to that determined by Sanger sequencing. To understand the influence of WGS depth on the accuracy of endogenous HHV-6 reconstruction, we downsampled the dataset to approximately 30x, 20x, 15x, 10x, and 5x autosome depths using the ‵picard DownsampleSam‵ function. Our pipeline detected endogenous HHV-6 at all read depths, and had accuracy near that of Sanger sequencing when the depth was higher than 15x. This demonstrates that, from moderate-to high-depth WGS datasets, our pipeline can reconstruct relatively accurate endogenous HHV-6 sequences suitable for phylogenetic analysis.

### Visualization of virus-chromosome hybrid reads

WGS reads that failed to map to the reference human genome were mapped to viruses using the pipeline described above. To detect virus-chromosome hybrid reads, read pairs with one read mapped to a virus and the paired read not mapped to the same virus were retrieved using ‵samtools view -f 8‵ command. Then, the unmapped mate reads were mapped to the reference human genome GRCh38DH using ‵blastn -evalue 1e-15 -culling_limit 2 - qcov_hsp_perc 90 -perc_identity 95 -word_size 11‵ command. Then, reads uniquely mapped to the human genome were retrieved and the mapped positions of the hybrid reads on the virus genomes and the human genome were visualized using a custom Python script. Because both SMRV and HTLV-1 have LTRs, reads mapped to 3’LTR were re-mapped to 5’ LTR. The script used for visualization is available from the following GitHub repository: ‘https://github.com/GenomeImmunobiology/Kojima_et_al_2020’.

### Phylogenetic analysis of endogenous HHV-6

To reconstruct phylogenetic trees of U regions, we used full-length genomes reconstructed by the endogenous HHV-6 reconstruction pipeline described above. The reconstructed sequences were aligned with known endogenous and exogenous HHV-6 using ‵mafft --auto‵ command. To exclude the regions thought to have low reconstruction accuracy, we removed DR and repeat sequences annotated in NCBI from the alignment. We removed regions corresponding to nucleotides 0-8089, 127548-128233, 131076-131854, 140075-140951, and 151288-159378 of HHV-6A NC_001664.4 and 0-8793, 9314-9510, 129045-129681, 133500-133863, 133981-134076, 140081-142691, and 153321-162114 of HHV-6B NC_000898.1 (all start positions here are 0-based numbering and end positions are 1-based numbering) using a custom Python script. Then, phylogenetic trees were inferred by the maximum likelihood method with the complete deletion option using MEGA X software. The Kimura 2-parameter model was used. The reliability of each internal branch was assessed by 100 bootstrap resamplings. The phylogenetic trees were visualized using ETEtoolkit.

To reconstruct phylogenetic trees of DR regions, we used DR reconstructed by the endogenous HHV-6 reconstruction pipeline described above. The reconstructed sequences are aligned with known endogenous and exogenous HHV-6 using ‵mafft --auto‵ command. To exclude the regions thought to have low reconstruction accuracy, we removed simple repeats and low complexity sequences from the alignment. To define simple repeats and low complexity sequences, we used RepeatMasker. We masked the reference HHV-6 (NC_001664.4 and NC_000898.1) with a ‵repeatmasker -s -no_is‵ command. We removed the regions corresponding to nucleotides 0-376, 1682-1730, 2302-2367, 2369-2451, 2692-2733, 3149-3181, 3433-3502, 3626-3670, 7483-7519, 7655-8089 of HHV-6A NC_001664.4 and 0-393, 1926-2011, 2674-2717, 3013-3067, 3670-3713, 3959-3988, 8248-8793 of HHV-6B NC_000898.1 (all start positions here are 0-based numbering and end positions are 1-based numbering) using a custom Python script. Then, phylogenetic trees were inferred and visualized as described above. The scripts used for phylogenetic analysis and the newick files of the phylogenetic trees are available from the following GitHub repository: ‘https://github.com/GenomeImmunobiology/Kojima_et_al_2020’.

### Processing of RNA-sequencing datasets

The SRA files of the Geuvadis RNA-seq dataset were downloaded from NCBI using the ‵prefetch‵ command in the NCBI SRA-tools. We used 159 datasets derived from LCL of Utah residents. The downloaded SRA files were converted to FASTQ files using ‵fasterq-dump‵ command in the NCBI SRA-tools with ‵-S‵ option. Paired-reads were then filtered using fastp software with ‵-l 20 -3 -W 4 -M 20 -t 1 -T 1 -x‵ options. The filtered paired-reads were then mapped to the human genome ‘GRCh38.p13.genome.fa’ downloaded from GenCode. For mapping, we used STAR software with ‵--quantMode GeneCounts --twopassMode Basic --outFilterType BySJout --outFilterMultimapNmax 20 --alignSJoverhangMin 8 -- alignSJDBoverhangMin 1 --outFilterMismatchNmax 999 --outFilterMismatchNoverReadLmax 0.04 --alignIntronMin 20 --alignIntronMax 1000000 --alignMatesGapMax 1000000‵ options. We provided the human gene annotation ‘gencode.v33.annotation.gtf’ downloaded from GenCode when indexing the reference human genome using STAR.

### Differential gene analysis

The 159 RNA-seq datasets were derived from 90 LCLs. Because some LCLs were represented by two different RNA-seq datasets, we merged the count tables originating from LCLs from the same donor. To remove low-expression genes from differential gene expression analysis (DE analysis), we calculated the average FPKM in 90 datasets and genes with FPKM lower than 1 were excluded from the downstream analysis. 43,585 genes were removed by this filtering, leaving 17,077 genes. 5 LCLs with a very low number of SMRV WGS reads (NA12286, NA12287, NA11930, NA12760, and NA11840), as these could potentially be derived from other SMRV-positive samples sequenced on the same lane, were excluded from the downstream analysis. The count tables generated by STAR were used for DE analysis. DE analysis was performed using the DESeq function in the DESeq2 package. For visualization, count tables were normalized by the counts function in the DESeq2 package with a ‵normalized=TRUE‵ option. Genes with p-value lower than 0.05 and changed by at least 2-fold were defined as genes with significant expression changes.

### *HERV-K* k*-mer counting*

If a presence/absence-type polymorphic HERV-K contains a unique region which distinguishes it from the other HERV-K loci, the presence/absence pattern of this HERV-K in humans should match to the presence/absence pattern of WGS reads originating from the unique region. To comprehensively find such *k*-mers, we exploited *k*-mer hashing of WGS reads. We first mapped WGS reads to a reference HERV-K (HERV-K113, NC_022518.1) and hashed mapped reads into *k*-mers. We excluded the LTR region from this analysis, because the LTR of HERV-K has a high similarity to SVA. Because of this exclusion criteria, our method captures *k*-mers derived from the HERV-K provirus, but does not detect polymorphisms of solo-LTRs.

FASTQ files of 2,504 1kGP high-coverage samples were downloaded from the following URL: ‘ftp://ftp.1000genomes.ebi.ac.uk/vol1/ftp/phase3/data/’. FASTQ files of the Han Chinese trio were downloaded from the following URL: ‘https://www.internationalgenome.org/data-portal/data-collection/structural-variation’. The FASTQ reads were mapped to the HERV-K reference sequence using Hisat2 with ‵--mp 2,1 --no-spliced-alignment‵ options and stored as BAM files. To exclude LTR regions for analysis, the reads mapped to 968-8504 of the HERV-K113 genome (start position is 0-based numbering and end position is 1-based numbering) were used for downstream analysis. To reduce the computational burden of *k*-mer counting, the HERV-K113 genome was split into 50-bp bins with a 10-bp sliding window. Then, sequences of the mapped reads corresponding to the HERV-K 50-bp bins were listed from the BAM files using a custom Python script. We defined those sequences as HERV-K *k*-mers. This detected 460,491 different HERV-K *k*-mers from 2,504 subjects in 1kGP. The occurrence of each HERV-K *k*-mer was counted by each sample using a custom Python script. The workflow described here was compiled as a Python pipeline and available from the following GitHub repository: ‘https://github.com/GenomeImmunobiology/Kojima_et_al_2020’.

To normalize the *k*-mer occurrence by the depth of human autosomes, we calculated the chromosome depths using ‵samtools coverage‵ command. For this calculation, we used CRAM files of the 2,504 high-coverage WGS provided from 1kGP (downloaded as described in the ‘*WGS datasets*’ section). Then, we calculated the mean depth of chromosome 1 to 22, which we refer to as the autosome depth. The *k*-mer occurrence of each dataset was divided by the calculated autosome and used as a normalized value.

### *Definition of high-frequency presence/absence-type HERV-K k*-mer*s*

To perform GWA analysis using polymorphic HERV-K *k*-mers, we decided to focus on HERV-K *k*-mers above a certain frequency threshold. Very rare *k*-mers would often be individual- or population-specific and thus be less informative for findingg association with SNVs from the trans-ethnic datasets. Therefore, we discarded presence/absence-type *k*-mers which were detected in less than 50 subjects (n=8,642) and we defined the remaining ones as high-frequency HERV-K *k*-mers.

### *Clustering of HERV-K k*-mer*s*

To perform hierarchical clustering of the frequencies of the presence/absence-type HERV-K *k*-mers by the 26 human populations, the mean *k*-mer frequencies in each population was first calculated. Then the mean *k*-mer frequencies of the 26 populations were clustered with Ward’s method in the clutermap function in the seaborn Python package.

To perform hierarchical clustering of the Pearson correlation coefficient of the high-frequency presence/absence-type HERV-K *k*-mers, we generated an all-by-all Pearson correlation coefficient matrix for presence/absence patterns of *k*-mers and performed hierarchical clustering of *k*-mers. The Pearson correlation coefficient was calculated by the corr function in the Python Pandas package. The *k*-mers were then clustered by Ward’s method in the clutermap function in the seaborn Python package.

Prior to GWA analysis, we formally defined clusters using DBSCAN. We clustered the all-by-all Pearson correlation coefficient matrix for presence/absence patterns of *k*-mers. We used the DBSCAN function in the Python scikit-learn package. Any mutation in the HERV-K provirus should actually result in 5 different *k*-mers, because we listed up *k*-mers corresponding to the reference HERV-K sequences scanned by 50-bp bins with a 10-bp window. Therefore, we used 5 for the ‵min_samples‵ parameter. We used 2.5 for the ‵eps‵ parameter. To determine appropriate epsilon, we performed DBSCAN using 0.5, 1.0, 1.5, 2.0, 2.5, 3.0, 3.5, and 4.0. The clustering results were visually cross-referenced with the result of hierarchical clustering, and an epsilon of 2.5 was chosen because it showed good concordance with the results of the hierarchical clustering while defining few enough clusters to allow GWA analysis using a reasonable computational load. The largest cluster defined by DBSCAN contained 175 unique *k*-mers.

### GWA analysis

The presence and absence of HERV-K *k*-mers in *k*-mer clusters defined by DBSCAN were then converted to categorical values. To reduce the computational cost of GWA analysis, we generated a consensus presence/absence pattern of *k*-mers in each cluster defined by DBSCAN. If a WGS dataset contained more than 80% of the *k*-mers in a *k*-mer cluster, the dataset was considered as a *k*-mer-positive dataset and labeled as 1, while a WGS dataset contained no or 80% or less number of *k*-mers in the *k*-mer cluster, the dataset was considered as a *k*-mer-negative dataset and labeled as 0. These presence/absence binary categorical values were used for the GWA analysis. For SNV annotations, we used GRCh38_v1a downloaded from ‘ftp://ftp.1000genomes.ebi.ac.uk/vol1/ftp/data_collections/1000_genomes_project/release/20181203_biallelic_SNV/’. We first removed SNVs with low frequency (< 1%), those violating Hardy-Weinberg equilibrium (1e-05), and those with high missing call rate (> 5%) using plink2 software with ‵--geno 0.05 --hwe 0.00001 --maf 0.01‵ options. Then the SNVs were pruned using plink2 software with ‵--maf 0.05 --indep-pairwise 100kb 0.5‵ options. PCA was performed using plink2 software using pruned SNVs. Association analysis was performed using plink2 with covariates and binary categorical values representing the presence and absence status of *k*-mers. The sex of WGS datasets and the eigen vectors generated by PCA were used as covariates. Because we performed 599 association tests, we set 8.33e-11 as the genome-wide significant p-value threshold.

### *Detection of HERV-K k*-mer*-associated loci*

To determine genome loci associated with the HERV-K *k*-mer clusters in GWA analysis, we first defined genome regions with association by each *k*-mer cluster. If two SNVs with significant p-values were within 1 Mb of one another, we considered those two SNVs to be within the same *k*-mer-associated locus. Otherwise, we considered the two SNVs to be in two separate *k*-mer-associated loci. If a *k*-mer-associated locus harbored 10 or more SNVs, we considered the genome region as a *k*-mer-associated locus, and the largest continuous region containing SNVs with association p values below the 8.33e-11 threshold were defined as a *k*-mer-associated loci. We detected 589 loci from 503 *k*-mer clusters. Because the same locus was detected in multiple GWA analyses with different *k*-mer clusters, we merged 589 genome regions using BEDTools merge command. Finally, we obtained 79 genome loci associated with HERV-K *k*-mers. The HERV-K *k*-mer-associated loci are available from the following GitHub repository: ‘https://github.com/GenomeImmunobiology/Kojima_et_al_2020’.

### Evaluation of LDfred to detect unknown HERV-K polymorphisms

To assess the sensitivity of LDfred to detect previously unknown polymorphisms, we used structural variations (SVs) in three subjects (NA12878, NA19434, HG00268) called by Audano et al. We extracted deletions that intersect with HERV-K annotated by RepeatMasker using the repeat library version 24.01 from Repbase. We detected 24 deletions in at least one in the three subjects. These 24 deletions ranged from 72 to 9,468-bp. We checked if these HERV-K SVs were located within loci associated with *k*-mer clusters identified by LDfred. Seventeen out of 24 were located with loci associated with *k*-mer clusters. Next we checked the consistency of the presence-absence patterns between *k*-mers and the deletions. When the *k*-mer presence-absence pattern of a *k*-mer cluster and the presence-absence pattern of the overlapping deletions were exactly the same, we considered that the LDfred result accurately reflected the HERV-K polymorphism. For example, if a presence pattern of *k*-mers in a *k*-mer cluster is (NA12878 = +, NA19434 = -, HG00268 = +) and the presence of HERV-K in the associated locus has the same pattern (NA12878 = +, NA19434 = --, HG00268 = +), we considered that LDfred detected the HERV-K polymorphism. On the other hand, if a presence pattern of *k*-mers in a *k*-mer cluster was (NA12878 = +, NA19434 = -, HG00268 = +) and the presence of HERV-K in the associated locus has different pattern (NA12878 = -, NA19434 = -, HG00268 = +), we considered that the LDfred result did not accurately reflect the HERV-K polymorphism. We detected 9 loci associating with *k*-mer clusters which harbor polymorphic HERV-K with consistent presence-absence patterns.

### Dot matrix analysis

The PacBio alignments to the human genome were downloaded from the following URL: ‘http://ftp.1000genomes.ebi.ac.uk/vol1/ftp/data_collections/hgsv_sv_discovery/working/’. To generate a dot matrix between a PacBio sequence and the reference human genome, first, similar sequences between two input sequences were detected and aligned using blastn with ‵-evalue 1 -word_size 7 -dust no‵ options. Then, the alignment was visualized by a custom Python script. The script used for this analysis is available from the following GitHub repository: ‘https://github.com/GenomeImmunobiology/Kojima_et_al_2020’.

### Detection of SMRV by PCR

The following cell lines were obtained from the NIGMS Human Genetic Cell Repository at the Coriell Institute for Medical Research: GM18998, GM18999, GM12878, GM12399, and GM11920. To confirm the existence of SMRV DNA in LCLs, we designed SMRV-specific primers and amplified SMRV DNA by PCR. Total DNA extracted from GM12399, GM11920, GM18998 were used as PCR templates. PCR primers used are listed in the Supplementary Table 5.

### Amplification of HERV-K by PCR and mapping to the human genome

Genome regions containing HERV-K in interest were amplified by PCR. Total DNA extracted from GM18998, GM18999, GM12878 were used as PCR templates. The amplicons were barcoded and sequenced using an Oxford Nanopore flongle flow cell. We obtained 18,702, 30,651, and 18,604 reads from each subject which passed standard minKNOW v3.6.5 quality control from these LCLs, respectively. The reads were mapped to GRCh38DH by bwa mem with the ‵-Y -K 1000000 -x ont2d‵ option. Because the HERV-K sequences could potentially be mis-aligned to multiple HERV-K loci, reads harboring sequences which mapping to the non-HERV-K regions at the termini of each PCR amplicon were extracted using a custom script. Finally, we obtained 2,928, 6,676, and 5,660 mapped reads, respectively. PCR primers used are listed in the Supplementary Table 5. A mutation rate of 0.5 × 10^−9^ substitutions per base, per year was assumed to estimate the age of the novel haplotypes (41).

### Software versions

Python 3.7.4

scikit-learn 0.22.1

biopython 1.74

pandas 0.25.1

seaborn 0.10.1

pysam 0.15.2

MEGA X 10.0.5

MAFFT v7.407

ete3 3.1.2

STAR 2.7.3a

R 3.6.1

DESeq2 1.22.2

BLAST 2.9.0+

samtools 1.10

bedtools v2.29.2

bcftools 1.9

Hisat2 version 2.2.0

PLINK v2.00a2.3LM

prefetch 2.9.3

fasterq-dump 2.9.6

bamCoverage 3.4.1

RepeatMasker 4.0.9

## Acknowledgements

The authors wish to acknowledge the resources of 1,000 Genomes Project and HGDP-CEPH Human Genome Diversity Cell Line Panel. We thank Thomas Sasani, Lynn Jorde, Aaron Quinlan, Julie E. Feusier, and Cody Steely for providing unmapped reads from phs001872, and Mark Lathrop for helpful discussions about LCLs generated by CEPH. The super-computing resources were provided by Human Genome Center, the Institute of Medical Science, the University of Tokyo (SHIROKANE), and the Office for Information Systems and Cybersecurity, RIKEN (HOKUSAI General Use project G20021). N.F.P. acknowledges funding from Cluster for Pioneering Research under the Hakubi fellowship program and from the discretionary budget of the Director of the RIKEN Center for Integrative Medical Sciences, Dr. Kazuhiko Yamamoto.

---

**Supplementary Figure 1.**
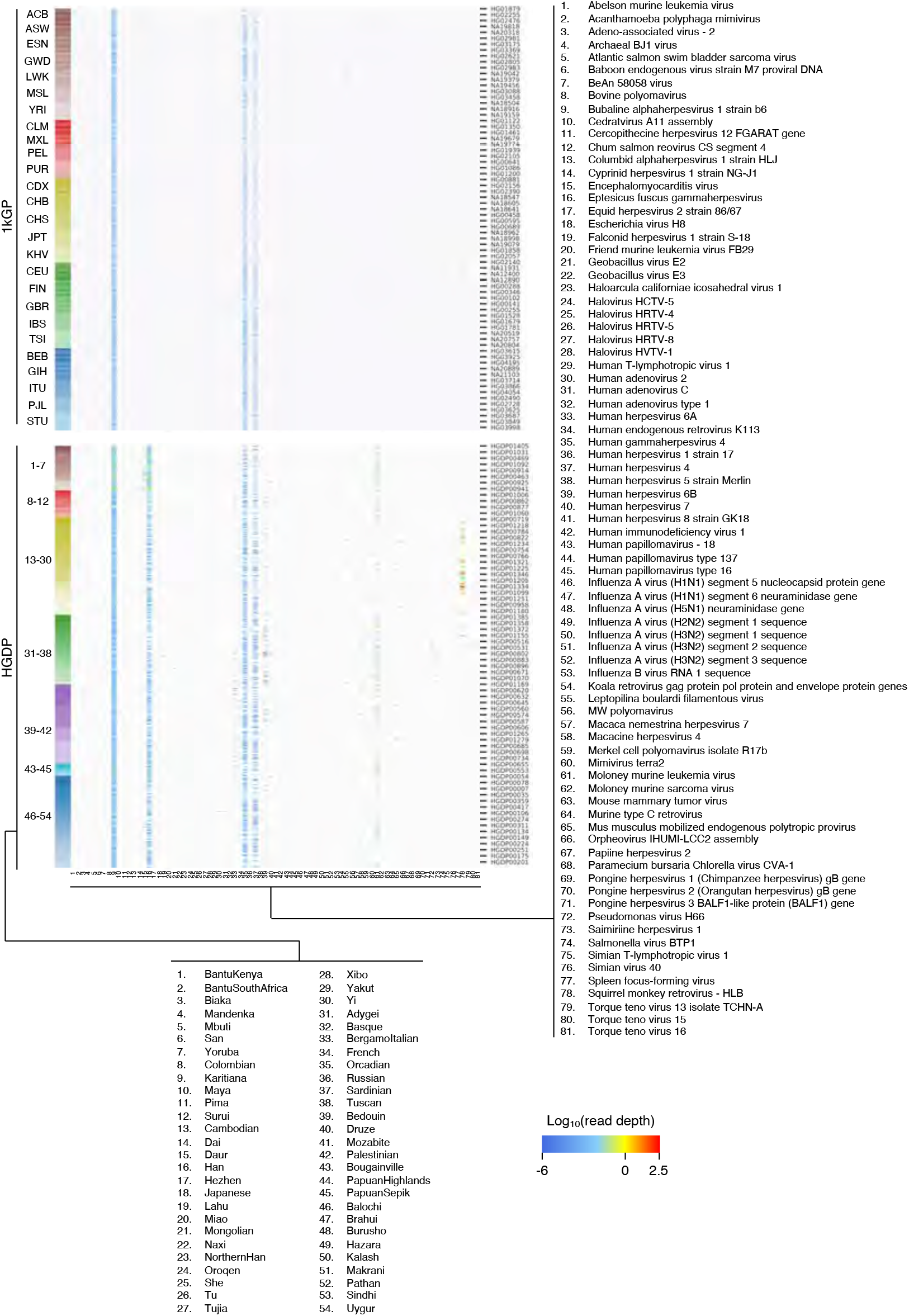
Virus search from 1kGP and HGDP WGS Heatmap shows the read depth of viruses with at least one read in at least one dataset from 2,504 1kGP as well as 808 HDGP datasets.

**Supplementary Figure 2.**
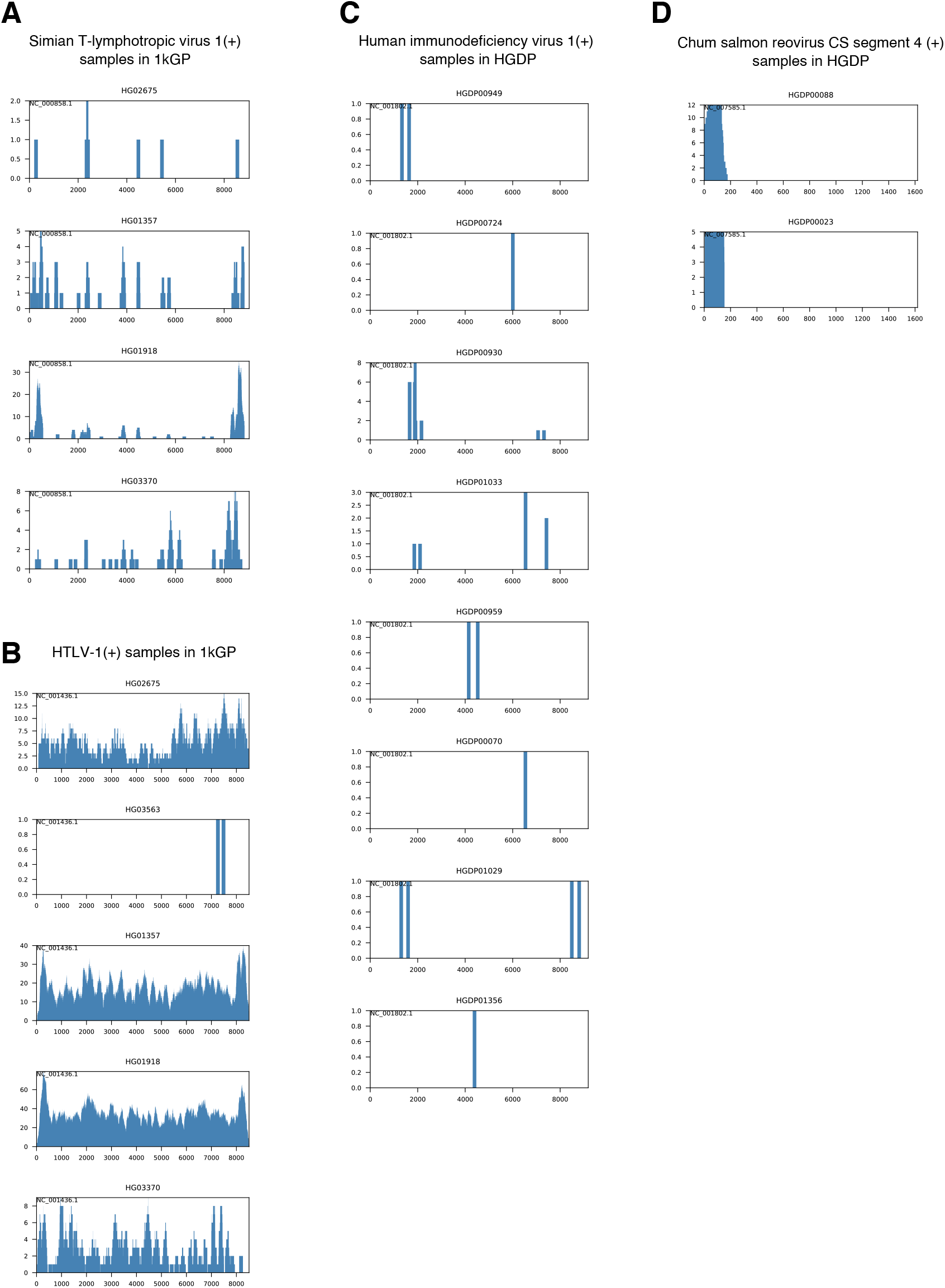
Read depth of STLV-1, HIV-1, and chum salmon reovirus Depth of reads mapping to STLV-1 (A), HTLV-1 (B), HIV-1 (C), and chum salmon reovirus (D) are shown. X-axis and Y-axis show the genome position of indicated virus and the depth of reads mapping to the indicated virus, respectively. The name of each dataset is shown at the top of the panel.

**Supplementary Figure 3.**
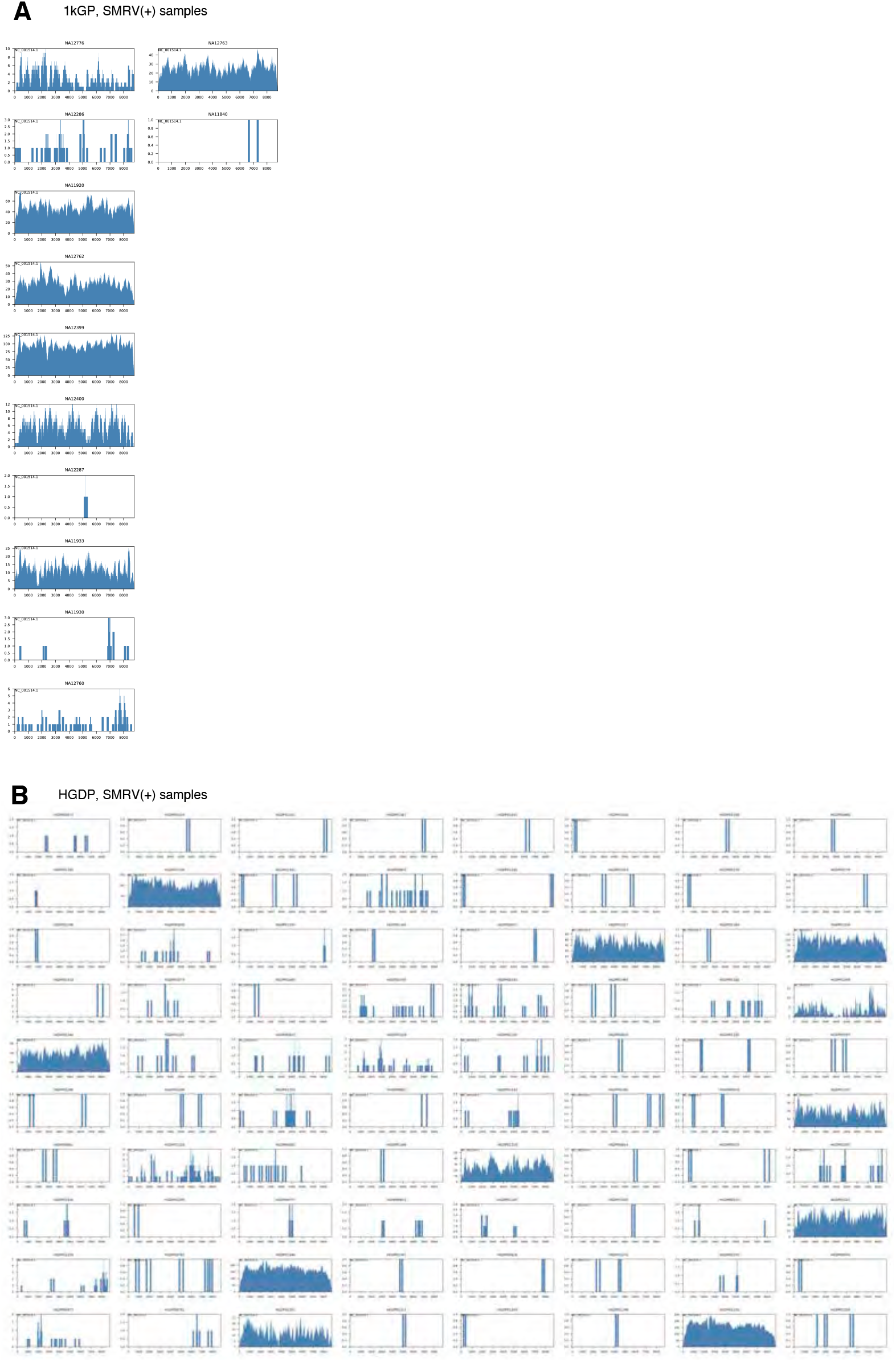
Read depth of SMRV and HTLV-1 Depth of reads from 1kGP (A) and HGDP (B) datasets mapping to SMRV-H are shown. X-axis and Y-axis show the genome position of indicated virus and the depth of reads mapping to the indicated virus, respectively. The name of each dataset is shown at the top of the panel.

**Supplementary Figure 4.**
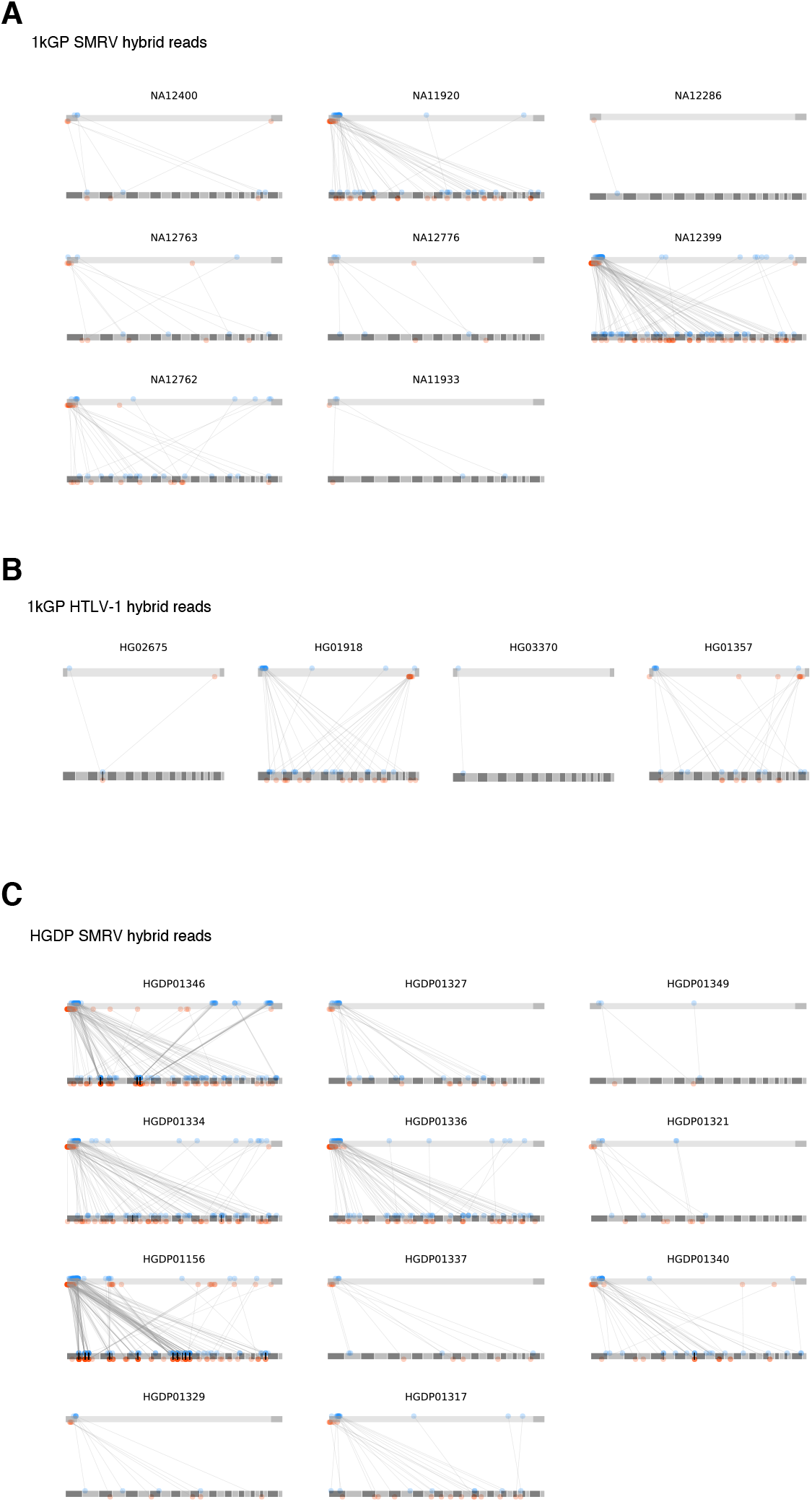
Virus-chromosome hybrid reads WGS read pairs which are mapped to both the virus genome and the human genome are shown. Gray bar in the top of each panel shows the SMRV-H (A), HTLV-1 (B), and SMRV-H (C). LTR are shown as dark gray rectangles. Gray bars in the bottom of each panel show the chromosome 1 to 22, X, and Y (from left to right). Read-1 and Read-2 of a read-pair are connected with a line. Reads mapping to forward and reverse directions are shown as blue and red dots, respectively. The reads mapping to left LTR was kept when a read was multi-mapped to both left and right LTR. The genome position of reads mapping only to right LTR were replaced to the left LTR. The name of each dataset is shown at the top of the panel.

**Supplementary Figure 5.**
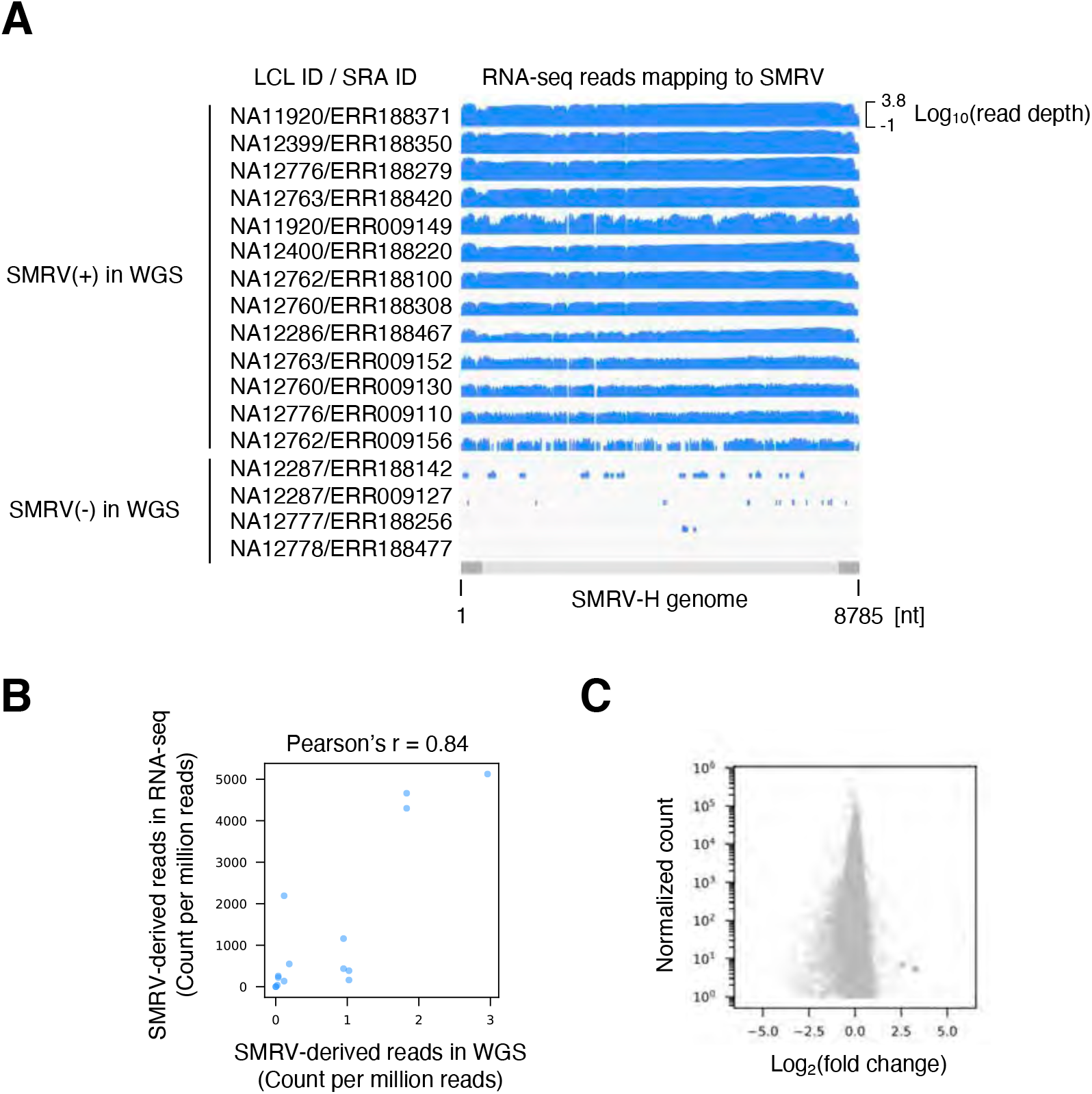
Differential gene analysis between SMRV-positive and -negative samples A. Depth of RNA-seq reads mapping to SMRV-H. Geuvadis RNA-seq datasets were used for the analysis. All datasets are shown with the same scale of the Y-axis. B. Correlation of the abundance of reads mapping to SMRV between WGS and RNA-seq. LCLs producing both WGS and RNA-seq were used for this analysis. C. MA plot showing the differences of gene expression and the normalized read counts. Two genes with differential expression are shown as blue dots.

**Supplementary Figure 6.**
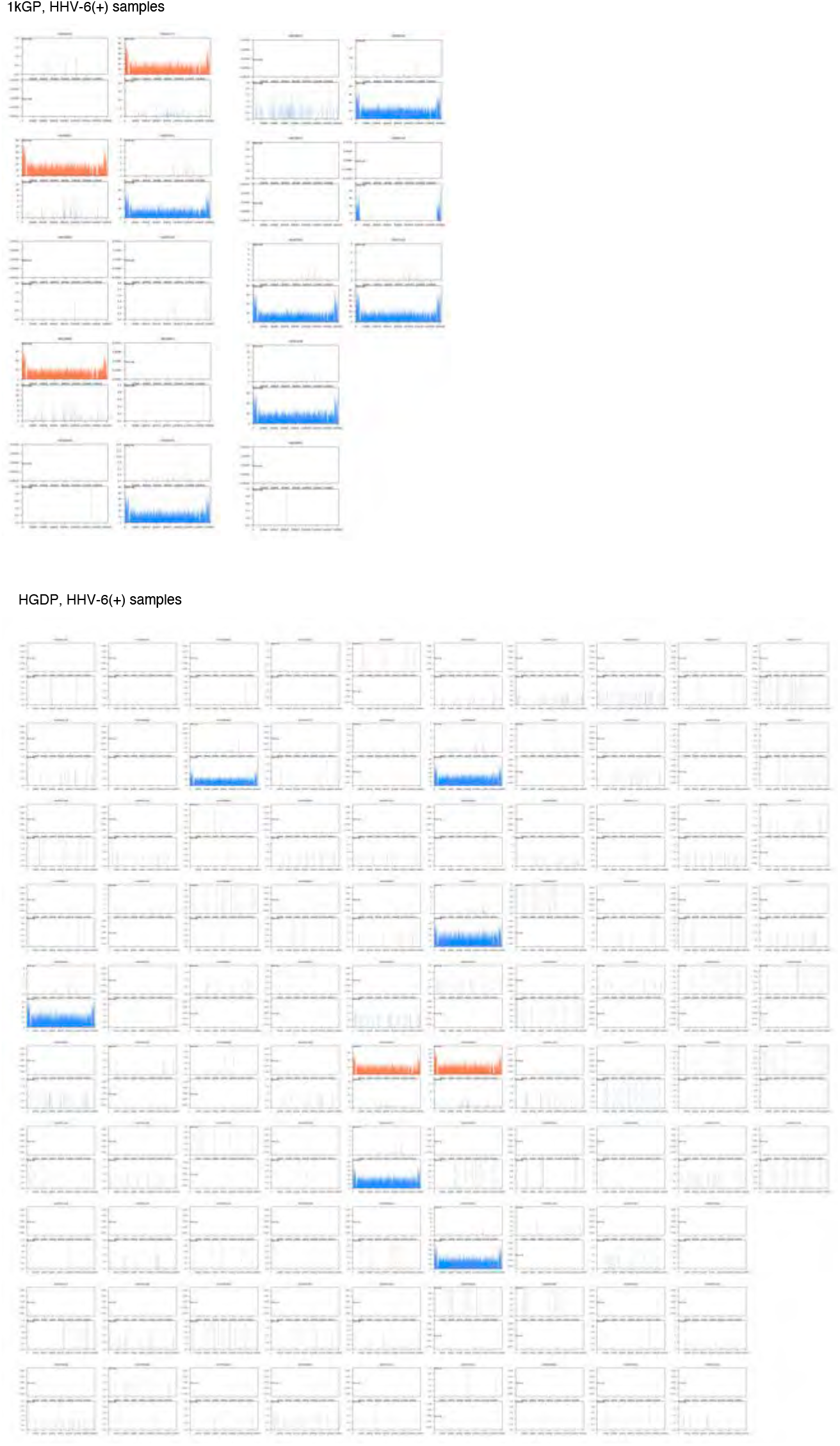
Read depth of HHV-6A and HHV-6B Depth of reads mapping to HHV-6A and HHV-6B are shown. Read depth of HHV-6A and HHV-6B are shown as orange and blue coverage plots, respectively. X-axis and Y-axis show the genome position of indicated virus and the depth of reads mapping to the indicated virus, respectively. The name of each dataset is shown at the top of the panel.

**Supplementary Figure 7.**
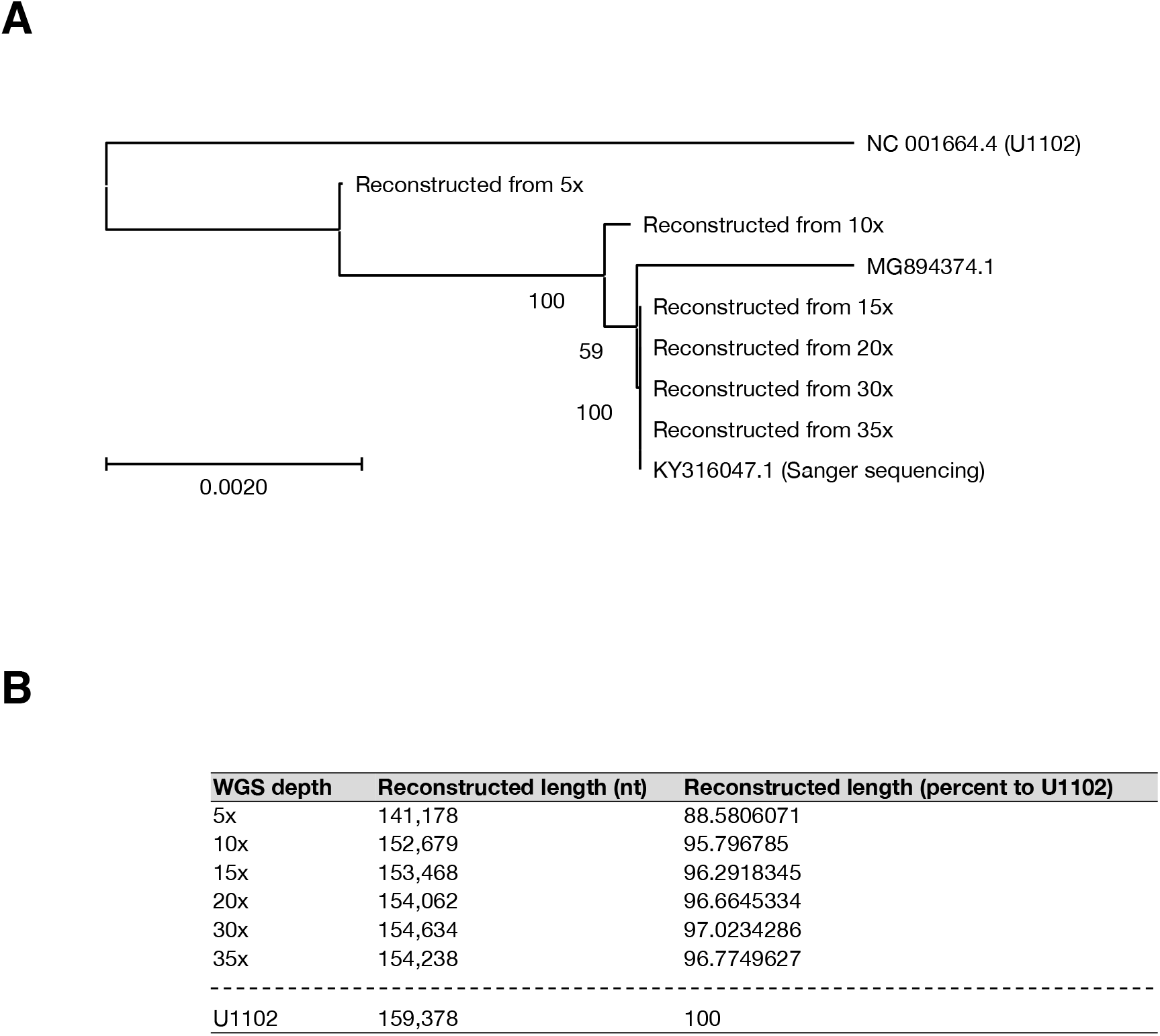
Accuracy of endogenous HHV-6 sequence reconstruction A. Phylogenetic analysis of reconstructed HHV-6A sequences. HHV-6A in NA18999 reconstructed from 35x, 30x, 20x, 15x, 10x, and 5x autosome depths are aligned with the reference HHV-6A (U1102) as well as the published sequences of HHV-6A Sanger sequenced using NA18999 DNA (KY316047.1) or reconstructed from a WGS of NA18999 (MG894374.1). B. The lengths of endogenous HHV-6A reconstructed from various WGS read depths.

**Supplementary Figure 8.**
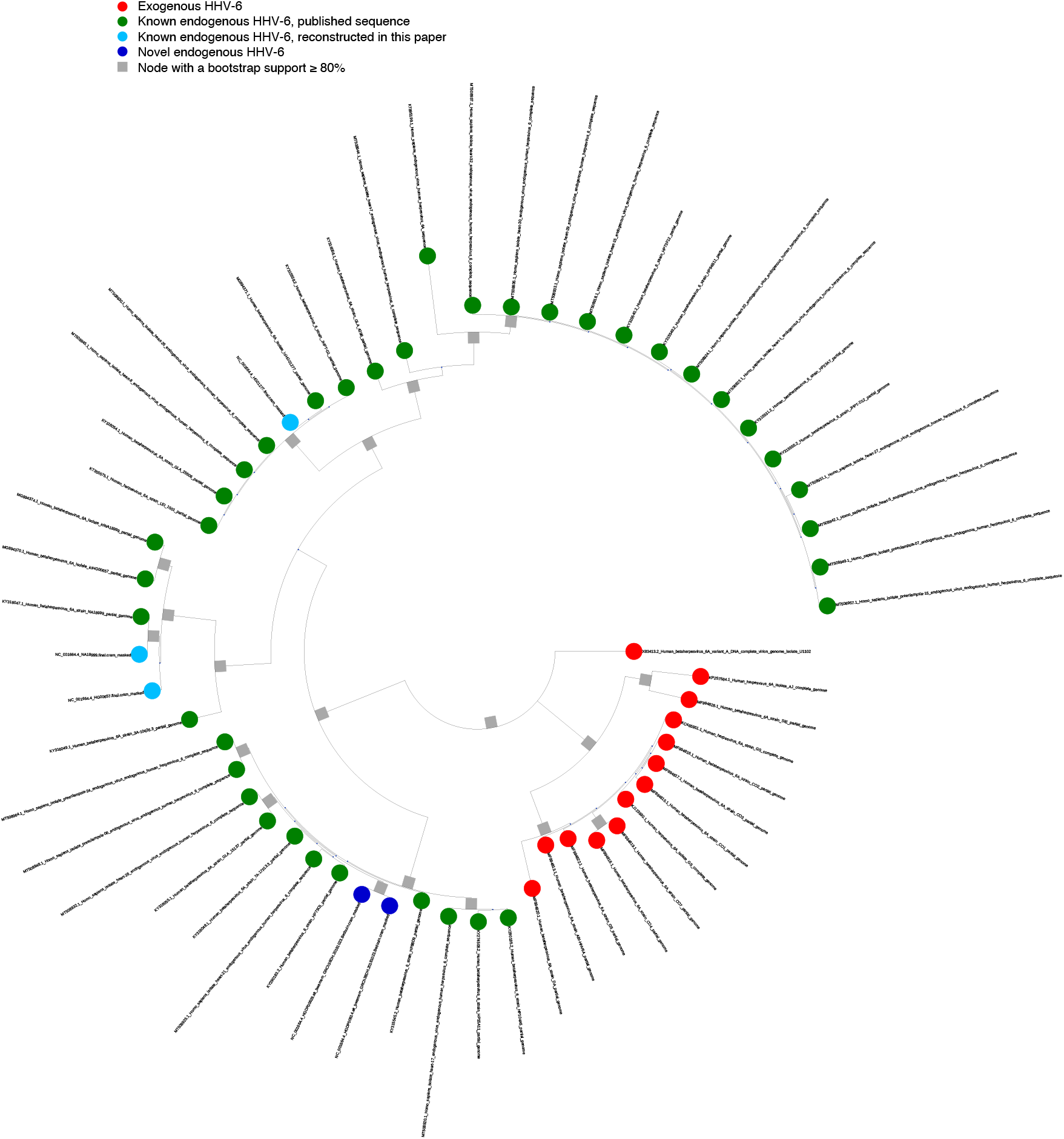
Phylogenetic tree of HHV-6A U with leaf names Phylogenetic trees inferred from U regions of HHV-6A. The publicly available sequences of endogenous and exogenous HHV-6A as well as ones reconstructed in the present study were used. The tree shown here is the same tree as Figure 3B left panel, except for showing the leaf names.

**Supplementary Figure 9.**
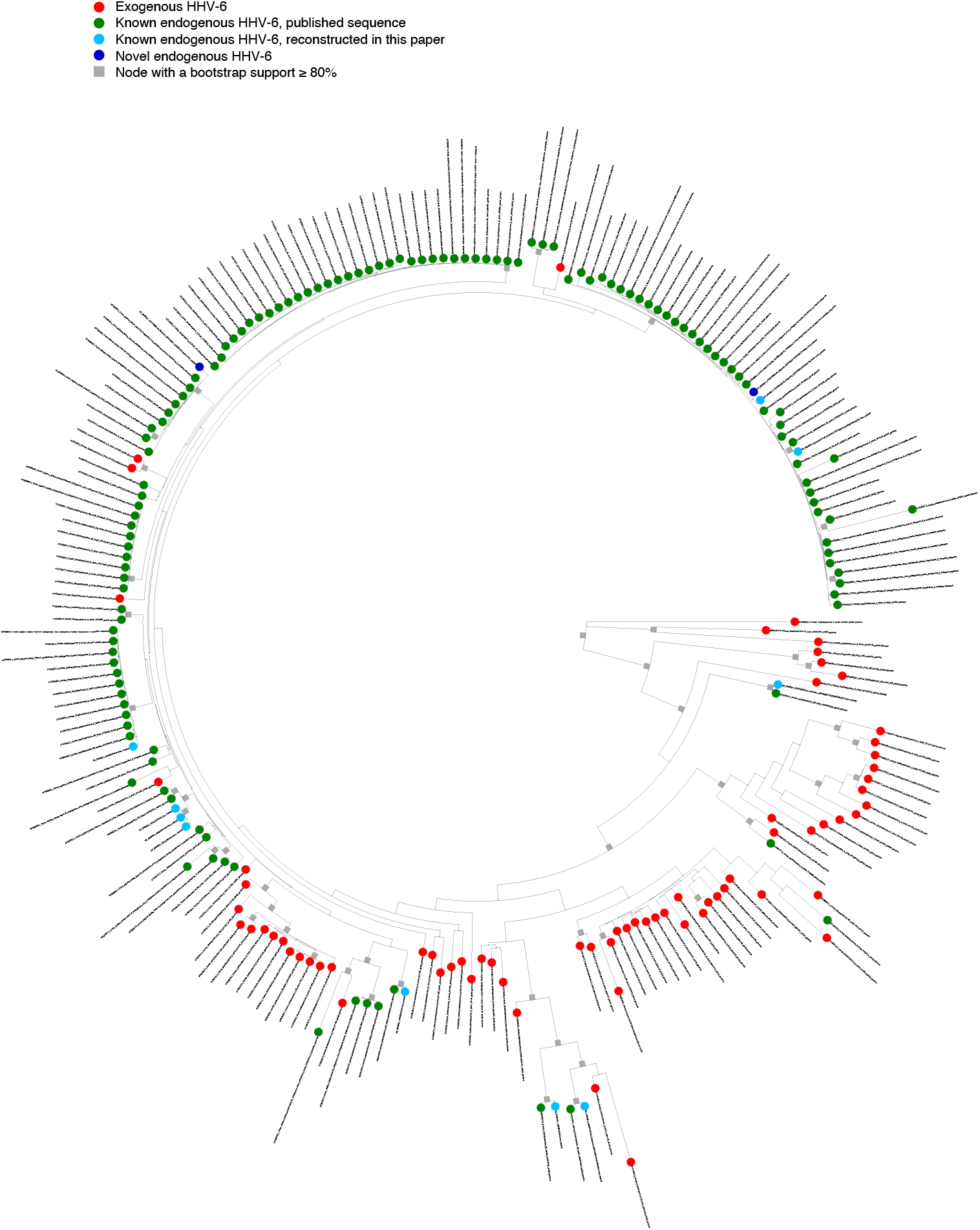
Phylogenetic tree of HHV-6B U with leaf names Phylogenetic trees inferred from U regions of HHV-6B. The publicly available sequences of endogenous and exogenous HHV-6B as well as ones reconstructed in the present study were used. The tree shown here is the same tree as Figure 3B right panel, except for showing the leaf names.

**Supplementary Figure 10.**
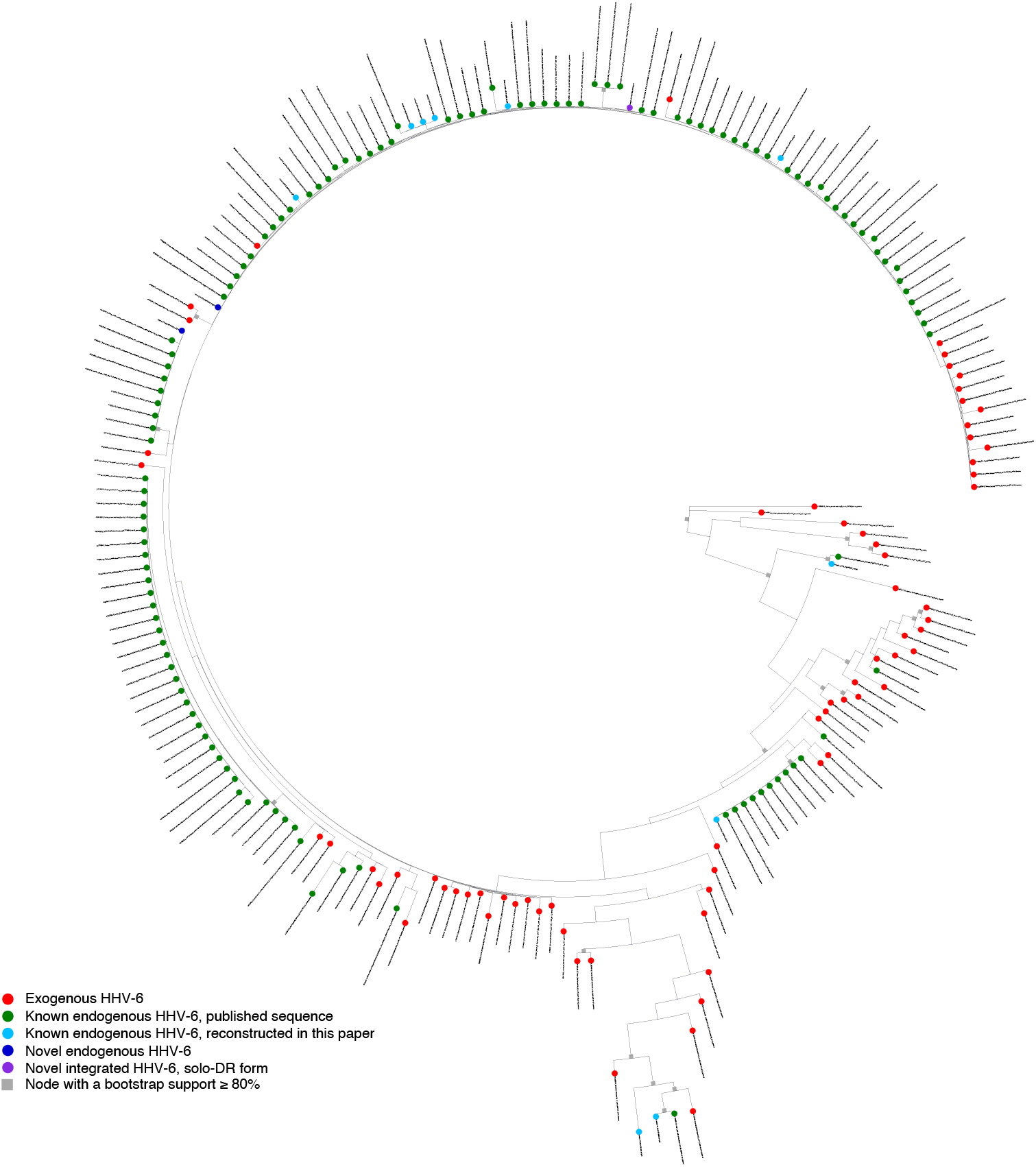
Phylogenetic tree of HHV-6B DR with leaf names Phylogenetic trees inferred from DR regions of HHV-6B. The publicly available sequences of endogenous and exogenous HHV-6B as well as ones reconstructed in the present study were used. The tree shown here is the same tree as Figure 3C, except for showing the leaf names.

**Supplementary Figure 11.**
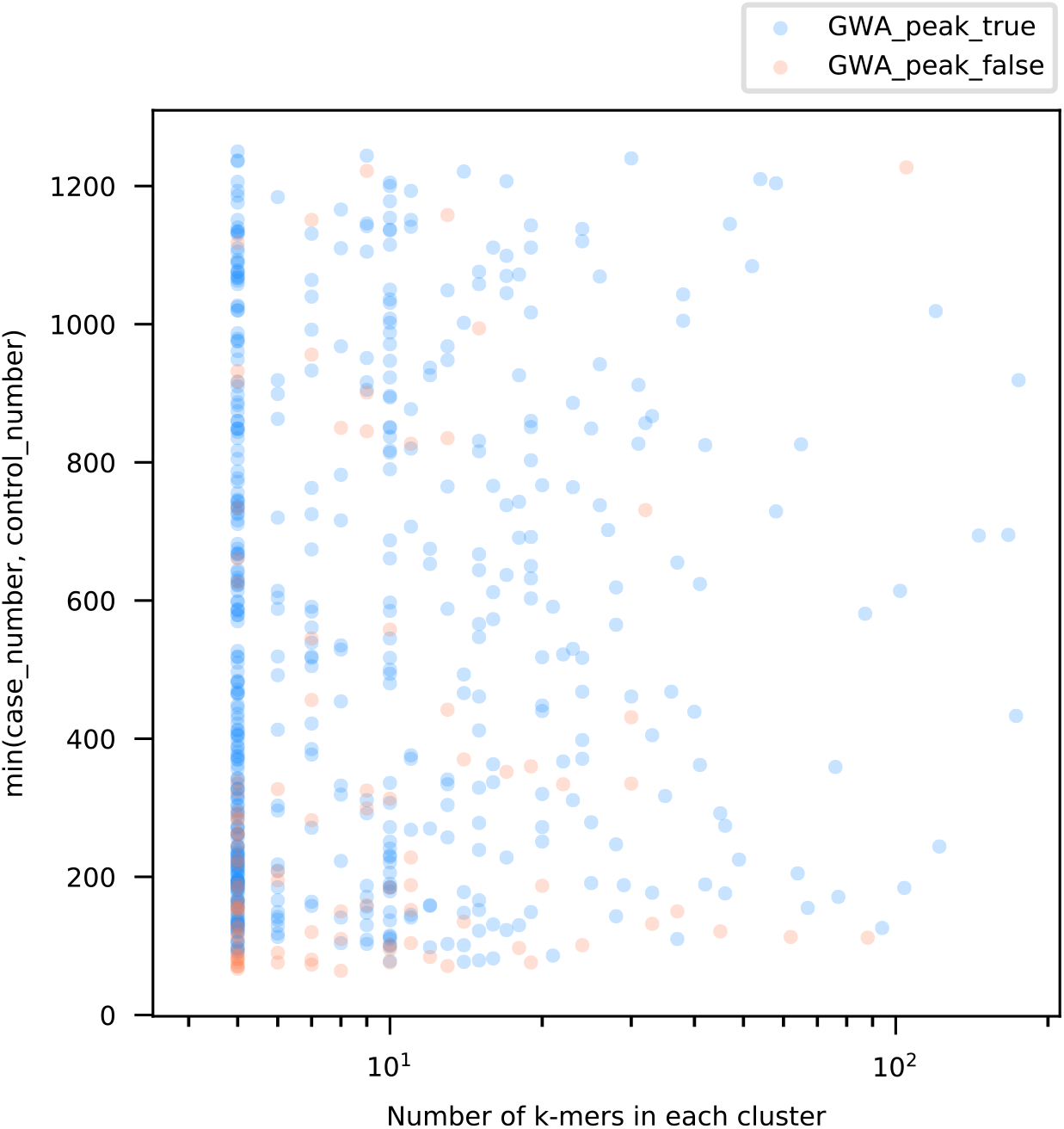
Numbers of *k*-mers in *k*-mer clusters Scatter plot shows the 597 *k*-mer clusters as dots. X-axis shows the number of *k*-mers in *k*-mer clusters. Y-axis shows the number of either case or control used for GWA analysis, whichever is smaller. Blue dots represent *k*-mer clusters with SNVs with association, while red dots show ones without any association to SNVs.

**Supplementary Figure 12.**
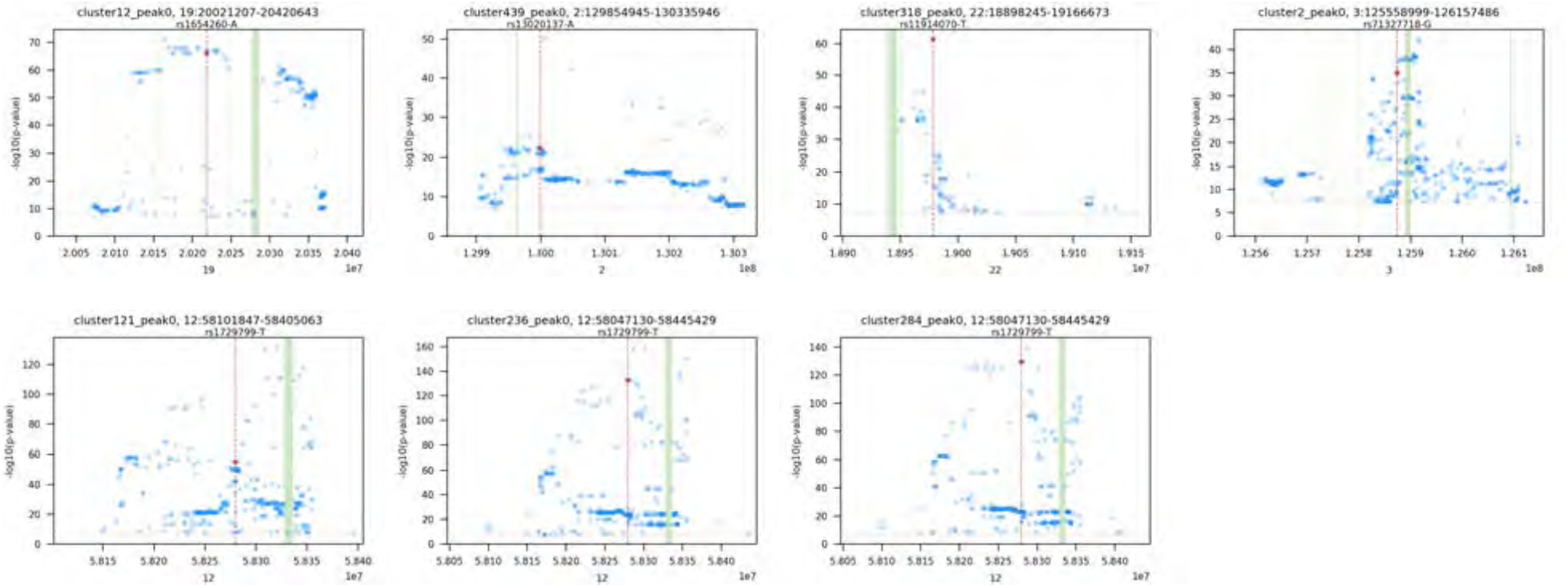
SNVs in the NHGRI-EBI GWAS catalog overlapping with the HERV-K *k*-mer LD regions Manhattan plots showing SNVs in the GWAS catalog overlapping with the indicated *k*-mer clusters. SNVs with p-value lower than 5e-08 are shown. Green lines show the reference HERV-K provirus. Red dots show the lead SNVs listed in the NHGRI-EBI GWAS catalog.

**Supplementary Figure 13.**
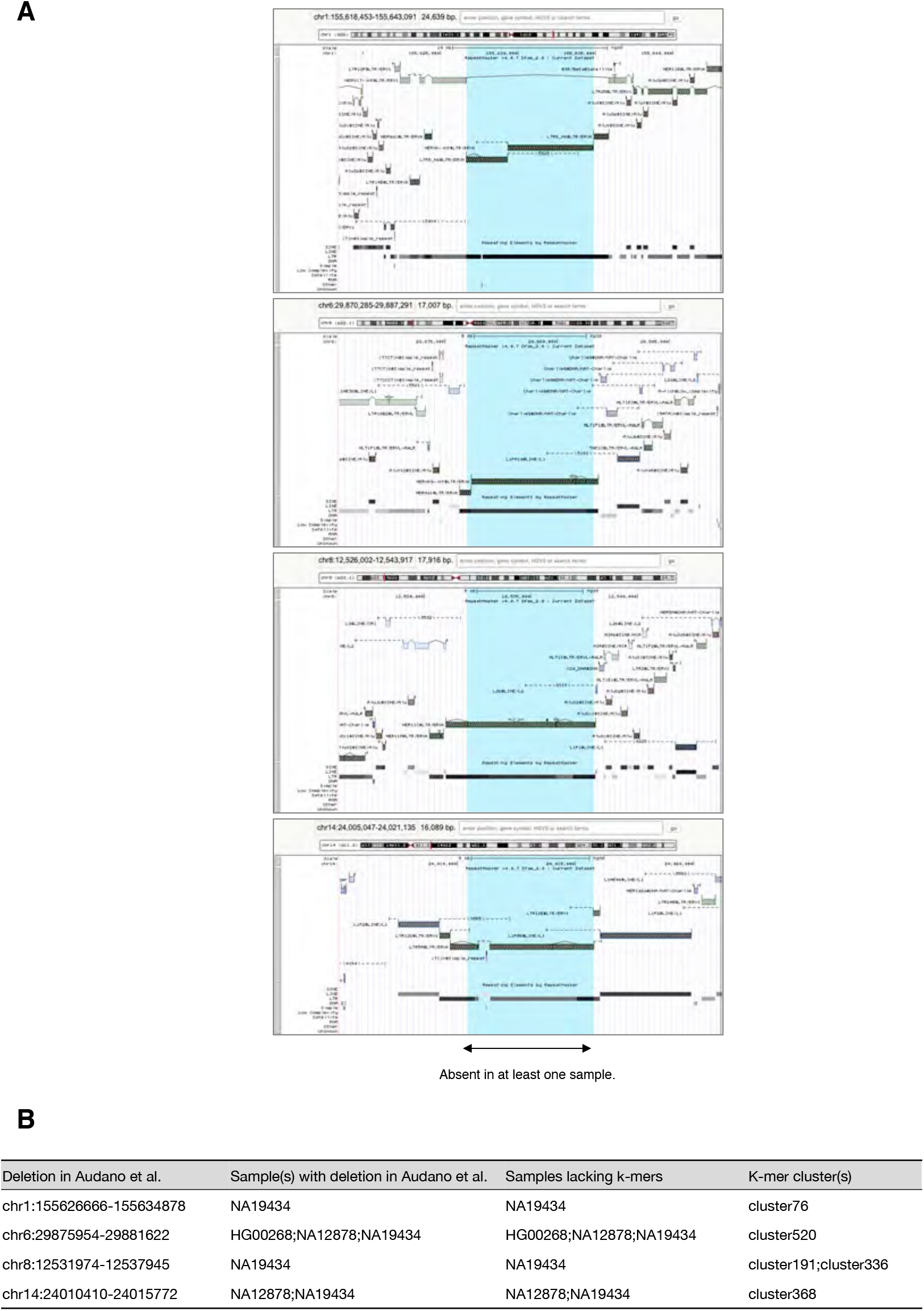
Provirus/solo-LTR-type HERV-K polymorphism captured by LDfred A. UCSC genome browser view showing Four polymorphic HERV-K. Blue regions are detected as sequence deletions in Audano et al. B. Cross-reference between deletions exist within HERV-K detected in Audano et al. and *k*-mer clusters detected by LDfred.

**Supplementary Figure 14.**
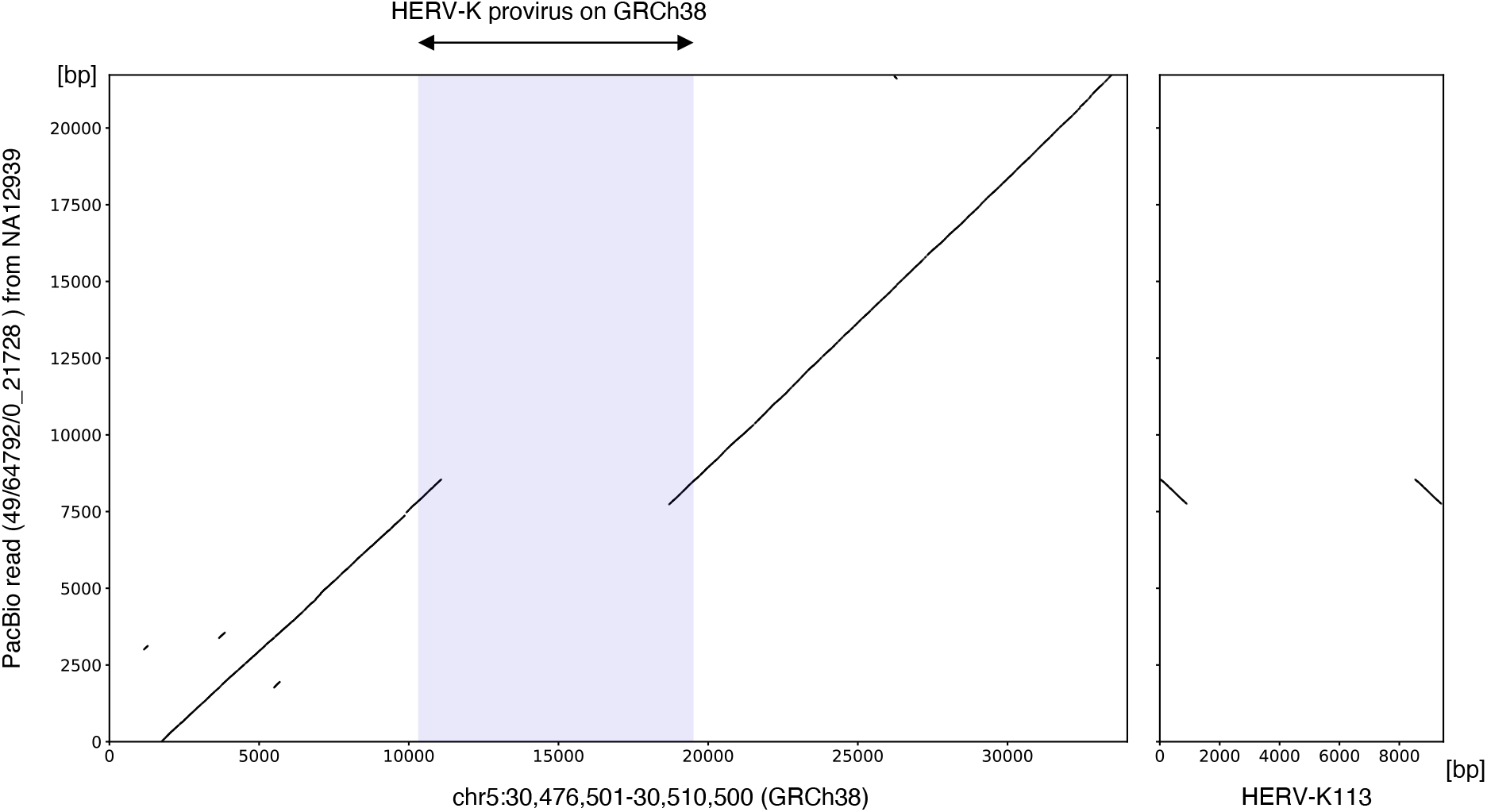
Provirus/solo-LTR-type HERV-K polymorphism captured by LDfred A PacBio read showing the absence of a HERV-K on chromosome 5 in NA12939. Left dot matrix shows the alignment between the partial sequence of chromosome 5 and PacBio read from NA12939 sequenced in Chaisson et al. Right dot matrix shows the alignment between HERV-K113 and the PacBio read. Blue region shows the provirus on chromosome 5.

**Supplementary Figure 15.**
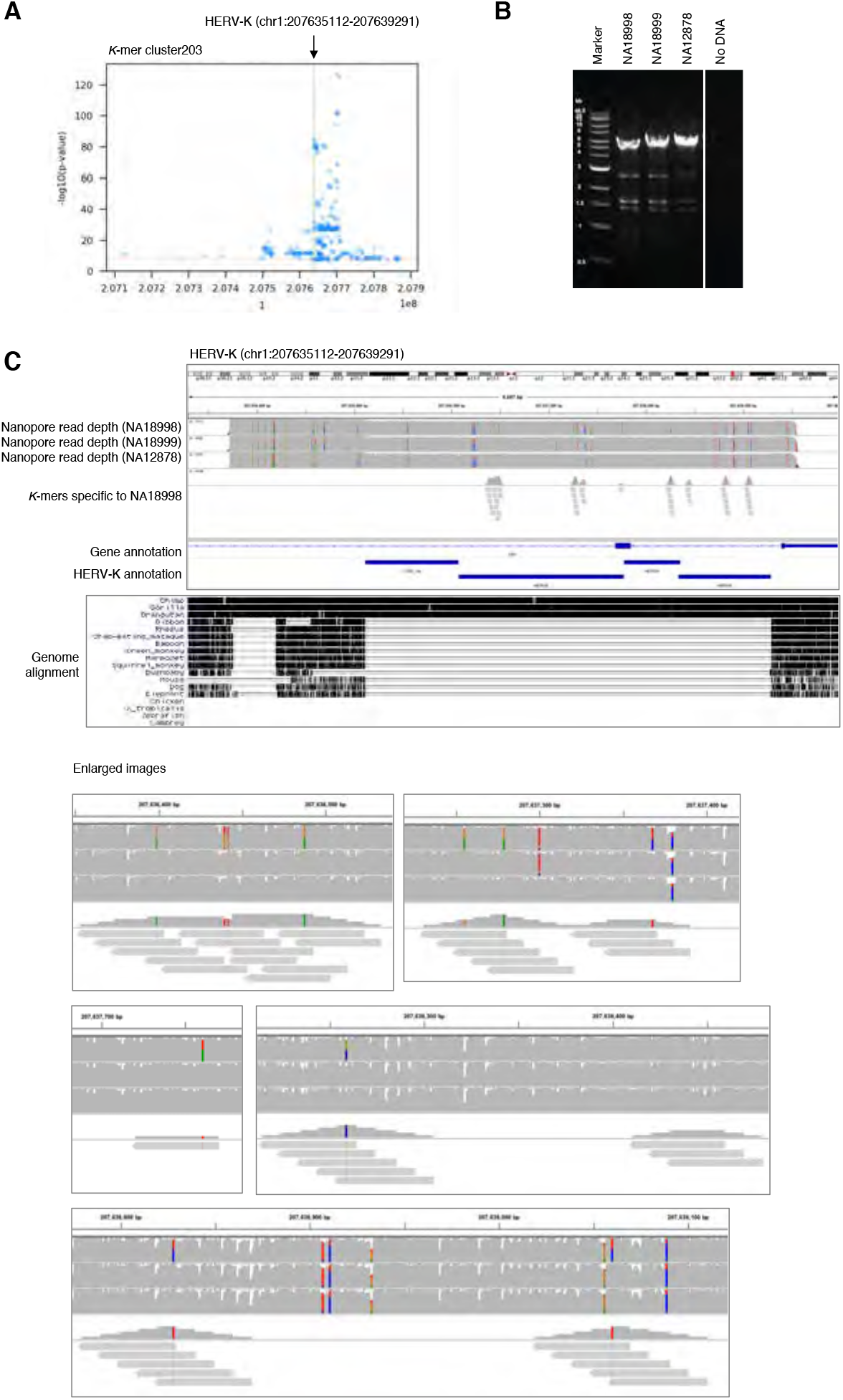
HERV-K SNVs captured by LDfred A. Manhattan plot showing SNVs associating the *k*-mer cluster203. SNVs with p-value lower than 5e-08 are shown. Green line shows the reference HERV-K provirus. B. Amplification of the HERV-K provirus by PCR. HERV-K provirus with adjacent sequence was amplified and PCR products were separated by gel electrophoresis. DNA extracted from LCLs originating from NA18998, NA18999, and NA12878 were used as templates. C. Upper panel: IGV view of long-read sequencing reads mapping to HERV-K. The PCR amplicons were sequenced using an Oxford Nanopore flongle flow cell and mapped to GRCh38. *k*-mers in *k*-mer detecting the HERV-K were also mapped to the PCR target regions. Lower panel: UCSC genome browser view showing the Multiz Alignment of 100 Vertebrates track.

**Supplementary Figure 16.**
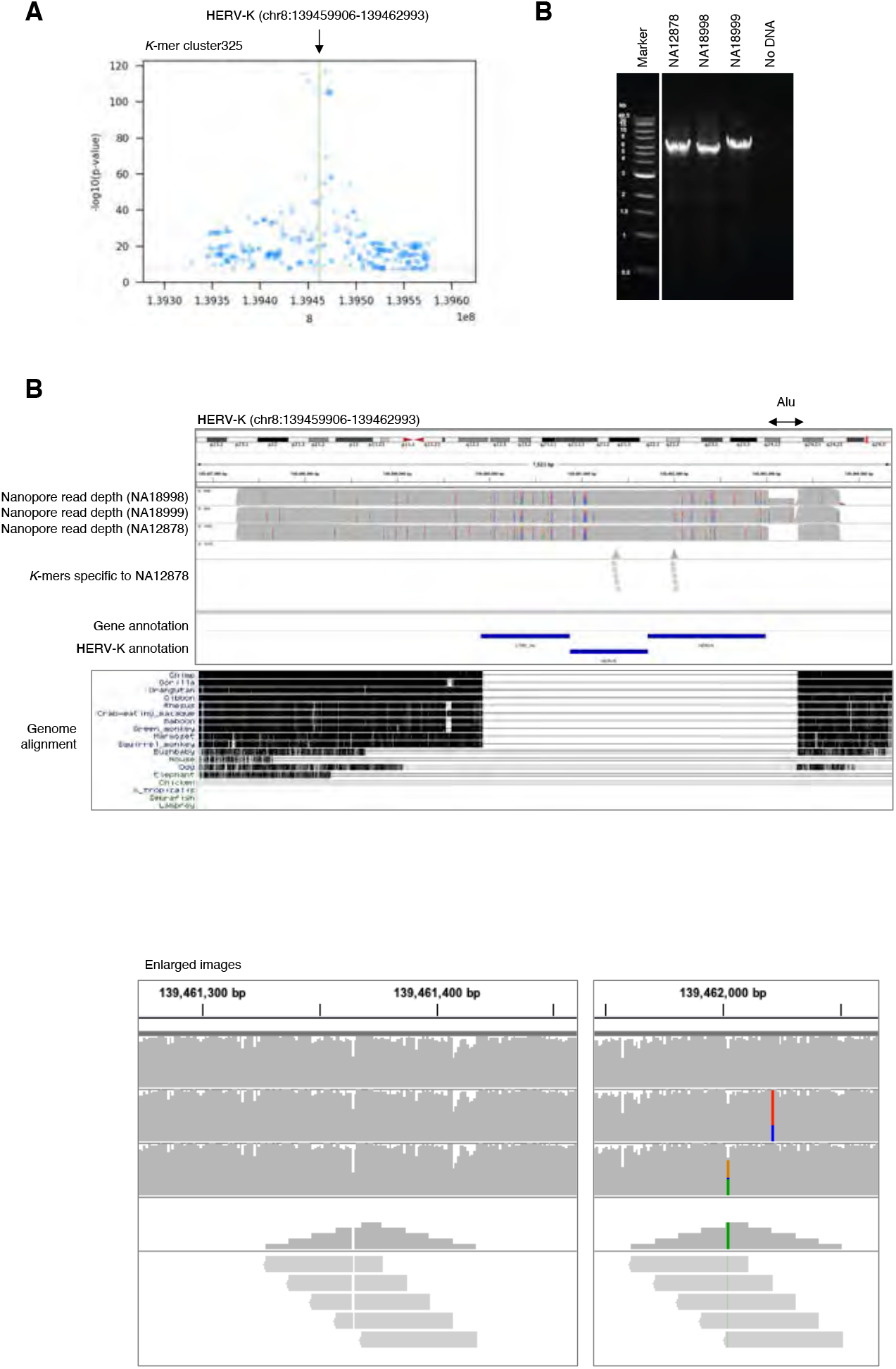
HERV-K SNVs captured by LDfred A. Manhattan plot showing SNVs associating the *k*-mer cluster325. SNVs with p-value lower than 5e-08 are shown. Green line shows the reference HERV-K provirus B. Amplification of the HERV-K provirus by PCR. HERV-K provirus with adjacent sequence was amplified and PCR products were separated by gel electrophoresis. DNA extracted from LCLs originating from NA18998, NA18999, and NA12878 were used as templates. C. Upper panel: IGV view of long-read sequencing reads mapping to HERV-K. The PCR amplicons were sequenced using an Oxford Nanopore flongle flow cell and mapped to GRCh38. *k*-mers in *k*-mer detecting the HERV-K were also mapped to the PCR target regions. Lower panel: UCSC genome browser view showing the Multiz Alignment of 100 Vertebrates track.

